# Increased presynaptic excitability in a migraine with aura mutation

**DOI:** 10.1101/719039

**Authors:** Pratyush Suryavanshi, Punam Sawant-Pokam, Sarah Clair, KC Brennan

## Abstract

Migraine is a common and disabling neurological disorder. The headache and sensory amplifications that characterize migraine are attributed to hyperexcitable sensory circuits, but a detailed understanding remains elusive. This is partly due to the paucity of genetic animal models associated with the common, less severe form of the disease. A mutation in casein kinase 1 delta (CK1δ) was identified in familial migraine with aura and advanced sleep phase syndrome. Spreading depolarization (SD), the phenomenon that underlies migraine aura, is facilitated in mice carrying one of the mutations (CK1δ_T44A_). However, the mechanism of this susceptibility is not known. We used a combination of whole-cell electrophysiology and imaging, *in vivo* and in *ex vivo* brain slices, to understand the cellular and synaptic underpinnings of this apparent circuit excitability. We found that despite normal synaptic activity and hyperpolarized neuronal membrane potentials at rest, CK1δ_T44A_ neurons were more excitable upon repetitive stimulation compared to wild-type (WT) controls. This was due to reduced presynaptic adaptation to high-frequency stimuli at excitatory, but not inhibitory synapses. Reduced adaptation at glutamatergic CK1δ_T44A_ synapses was mediated by a calcium-dependent enhancement of the size of the readily releasable pool of synaptic vesicles. This caused an increase in the cumulative amplitude of excitatory currents, and a higher excitation-to-inhibition ratio during sustained activity, both *in vivo* and in brain slices. Fluorescence imaging revealed increased glutamate release in CK1δ_T44A_ compared to WT brain slices, corroborating the presynaptic gain of function observed with electrophysiology. Action potential bursts elicited in individual CK1δ_T44A_ neurons enhanced glutamatergic feedback excitation within local microcircuits, further amplifying firing frequencies. At a network level *in vivo*, CK1δ_T44A_ mice showed increased duration of up state activity, which is dependent on recurrent excitation. Finally, we demonstrated that SD susceptibility of CK1δ_T44A_ brain slices could be returned to WT levels with the same reductions in extracellular calcium that normalized presynaptic adaptation. Taken together, these findings show a stimulus-dependent presynaptic gain of function at glutamatergic synapses in a genetic model of migraine, that accounts for the increased SD susceptibility and may also explain the sensory amplifications that are associated with the disease.

## Introduction

Migraine affects about 12% of the world population and causes enormous disability, especially to women and to those in working and childbearing years.^1,2^ Yet the disease remains poorly understood at the cellular and circuit level. Although migraine is typically associated with headaches, multiple lines of evidence suggest it is better characterized as a disorder of multi-sensory gain.^3^ In addition to the headache, migraine attacks involve amplification of sensory percepts including light, sound, smell, touch, and enteroception (allodynia, photophobia, phonophobia, osmophobia, and nausea, respectively).^4,5^ Migraine sufferers also exhibit larger amplitude sensory-evoked responses, and reduced habituation to repeated sensory stimulation during attacks and inter-ictal periods.^6–8^ In a third of migraineurs, the attack is preceded by an aura, which is caused by a massive spreading depolarization (SD) of cortical tissue.^3,9^

The mechanisms underlying the aberrant sensory network activity in migraine are poorly understood. Animal models of monogenic forms of migraine offer unique mechanistic insight,^10^ as mutations in single genes, and resultant dysregulation of downstream molecular pathways, produce migraine phenotypes. Established monogenic mouse models harbor mutations found in familial hemiplegic migraine (FHM types 1 and 2) and exhibit an increased probability of release and impaired clearance at glutamatergic synapses, respectively.^11–14^ These convergent lines of evidence suggest that increased synaptic excitability is a potential unifying mechanism of the disease. However, these models represent rare and severe forms of migraine, in which the migraine attacks are accompanied by hemiplegia.^11^ It is thus of interest to determine whether such mechanisms apply more broadly.

Loss of function mutations in casein kinase 1 delta (CK1δ) were recently identified in two families exhibiting familial migraine with aura and advanced sleep phase syndrome.^15^ Unlike FHM mutations, the CK1δ mutations segregate with phenotypically common migraine with aura. Mice carrying one of the mutations (CK1δ_T44A_) exhibit increased sensitivity to tactile and thermal hyperalgesia elicited by the migraine trigger nitroglycerin (NTG), as well as increased susceptibility to SD,^15^ thus confirming that two key phenotypes of the disease are represented in the animal model. However, the CK1δ_T44A_ mutation affects a ubiquitous serine-threonine kinase with broad roles regulating normal cellular function.^16–19^ Thus, although the molecular moiety responsible for the migraine phenotypes in CK1δ_T44A_ mice is known, the underlying cellular and synaptic mechanisms remain elusive. Here, using whole-cell electrophysiology and imaging *in vivo* and in brain slices, we uncovered presynaptic mechanisms contributing to SD susceptibility in CK1δ_T44A_ mice.

## Materials and methods

### Animal handling and experimental design

All experiments were conducted in accordance with and approved by the Institutional Animal Care and Use Committee at the University of Utah. Experiments were conducted on 2 – 3-month-old male CK1δ_T44A_ and wild-type (WT) littermate mice (line 827)^15,20^ weighing between 20 and 30 g. Animals were housed in temperature-controlled rooms on a 12-hour light-dark cycle. The sample size was determined using data from preliminary experiments as well as previous reports, generally *n*=5-9 mice for *in vivo* and *n*=15-20 slices (7-10 mice) for *in vitro* experiments. For patch-clamp electrophysiology, recordings were obtained from 1 neuron per mouse (*in vivo*) or 1-2 neurons per slice (*in vitro*). Experimenters were blind to the animal genotypes in all experiments. Most comparisons were made across genotypes within the same condition or across conditions within the same animal or slice (paired or repeated measures).

### *In vitro* brain slice preparation

CK1δ_T44A_ and WT littermate mice were deeply anesthetized with 4% isoflurane, and the brain was removed for slice preparation. Coronal sections were cut in ice-cold dissection buffer (in mM; 220 Sucrose, 3 KCl, 10 MgSO_4_, 1.25 NaH_2_PO_4,_ 25 NaHCO_3,_ 25 D-glucose, 1.3 CaCl_2_), and sections containing somatosensory cortex^21^ were allowed to recover in a chamber containing normal artificial cerebrospinal fluid (ACSF: in mM; 125 NaCl, 3 KCl, 1.3 MgSO_4_, 1.25 NaH_2_PO_4,_ 25 NaHCO_3,_ 25 D-glucose, 1.3 CaCl_2_, and saturated with 95%O_2_ / 5%CO_2_) at 35°C. For electrophysiology experiments, the slices were transferred to a submerged chamber constantly supplied with ACSF (flow rate: 2.5 mL/min, saturated with 95%O_2_ / 5%CO_2_) also maintained at 35°C. In experiments manipulating extracellular calcium concentrations ([Ca^2+^]_e_), ACSF containing 0.65 mM CaCl_2_ and 1.95mM MgCl_2_ was used *after* slice incubation.

### *In vitro* whole-cell electrophysiology

All whole-cell patch-clamp recordings were obtained from regular spiking pyramidal neurons^21^ in layer 2/3 (L2/3) somatosensory cortex visualized using differential interference contrast (DIC) microscopy. Whole-cell patch-clamp recordings were obtained using glass microelectrodes (4-6 MΩ resistance, tip size of 3-4 μm). To record intrinsic membrane properties, patch electrodes were filled with intracellular solution containing 130 K-gluconate, 5.5 EGTA, 10 HEPES, 2 NaCl, 2 KCl, 0.5 CaCl_2_, 0.3 Na_2_GTP, 2 MgATP, 7 phosphocreatine (concentrations in mM, pH = 7.2, 289-292 mOsm/kg-H_2_O). Baseline membrane voltage (resting membrane potential, V_m_), as well as membrane voltage responses to 20pA current injection steps (from −100 to +400pA, 500ms duration), were recorded. Spontaneous excitatory and inhibitory postsynaptic currents (sE/IPSCs) were recorded using patch pipettes containing 130 CsMeSO_4_, 3 CsCl, 10 HEPES, 2 MgATP, 0.3 Na_2_GTP, 5 EGTA, 10 Phosphocreatine, 5 QX-314, 8 biocytin (Concentration in mM; pH 7.2, 291-294 mOsm/kg-H_2_O). Excitatory and inhibitory currents were isolated by clamping neuronal membrane potential to −70mV (near inhibitory reversal potential) and 10mV (near excitatory reversal potential) respectively. EPSCs and IPSCs were pharmacologically blocked by DNQX (50 µM) and picrotoxin (PTX, 20 µM) respectively, confirming the AMPA and GABA_A_ receptor-mediated nature of the respective currents.

### Tonic inhibitory current recording

To isolate tonic inhibitory currents, GABA_A_ receptor-mediated phasic IPSCs were suppressed using 20 µM PTX for at least 5 minutes.^22,23^ A histogram of 10,000 data points (5 sec at 2KHz) was generated at baseline and after PTX treatment. A gaussian filter was fitted to the part of the distribution not skewed by phasic IPSCs to obtain mean holding currents.^22^ The PTX-sensitive tonic current was measured by subtracting the mean holding currents following PTX treatment from the baseline.

### Stimulus train evoked E/IPSCs

For stimulus train experiments, bipolar stimulation electrodes were fabricated from pulled theta glass pipettes (0.5 to 1GΩ resistance, 4-7 µm tip diameter) and filled with saline. Chlorinated silver wires were inserted into each half of the theta glass, and connected to the two poles of the stimulus isolator. Distinct barrels in the L4 somatosensory cortex were identified using IR-DIC microscopy.^24^ Theta glass stimulation electrodes were placed at the bottom edge of the barrels. Postsynaptic L2/3 neurons from the same cortical column were patched for voltage-clamp recordings. Postsynaptic responses to 100 µs stimuli ranging from 1 to 10 mV intensity were recorded to establish input-output relationships. The range of stimulus intensity was restricted to evoke only column-specific monosynaptic EPSC (at −70mV) and di-synaptic IPSC (at 10mV) responses. Paired-pulse ratios (PPRs) were calculated as the ratio of amplitudes of the second and the first evoked response (EPSC2/1, Figure 4). Stimulus intensity sufficient to evoke ∼50% of the maximum responses was selected for the rest of the experiment. Trains of 10 or 30 stimuli were introduced at different frequencies (10/20/50 Hz) and resulting monosynaptic EPSC and di-synaptic IPSC responses were measured by clamping post-synaptic cell voltage at −70mV and 10mV respectively.^25^

### *In vivo* whole-cell electrophysiology

Mice (males; 2-3 months old) were anesthetized using urethane (0.75 g/kg; i.p.) supplemented with isoflurane (∼0.5 %). Body temperature was monitored and maintained at 35 - 37°C using a heating pad. Vital signs (HR: 470–540 bpm, SpO_2_: 92-98%, respiration: 120-140/min) were monitored (MouseStat, Kent Scientific) throughout the experiment, and maintained within a physiologically normal range. An Omega-shaped head bar was mounted on the skull, using glue and dental cement, and affixed with screws to a holding rod attached to the stage. A craniotomy 2 mm in diameter was made above the hind paw region of the somatosensory cortex (1 mm caudal to the bregma and 2 mm lateral to the midline), and filled with 1.5% agarose (in ACSF) to keep the cortical surface moist and dampen the movement associated with breathing.^21^ We used *in vivo* whole-cell techniques to record the membrane potential from layer 2/3 pyramidal neurons in the primary somatosensory cortex (S1 L2/3 neurons) and analyzed spontaneous up states in current-clamp mode at resting membrane potential (i.e., −65 to −70 mV). Patch electrodes with 4-6 MΩ resistance and longer taper were used (tip size of 3-4 μm).^21^

### 2-photon laser scanning microscopy

Acute slices containing the somatosensory cortex were prepared and incubated similarly to whole-cell electrophysiology experiments. After incubation, slices were placed in a submerged chamber and perfused with ACSF (2 to 2.5 mL/min), under a 2-photon microscope (Neurolabware with Hamamatsu H11901 and H10770B photomultiplier tubes; Cambridge Technology CRS8 8KHz resonant scanning mirror and 6215H galvanometer scanning mirror; Nikon 16x/0.8 NA objective; Coherent Cameleon Discovery dual laser system [tunable laser = 920 nm; fixed laser = 1040 nm; pulse width ≈100 fs]; Semrock emission filters [510/84 nm green & 607/70 nm red]; power ≤ 25 mW; 250 µm – 1.5 mm field of view (FOV) on its long axis with ∼1.15-1.73µm/pixel resolution at 2X digital zoom, Scanbox acquisition software). Glutamate transients were evoked using a stimulation paradigm similar to the whole-cell electrophysiology experiments, in the presence of glutamate receptor antagonists (CNQX 20µM, AP5 50µM).^26^ Maps of glutamate transients across layers were acquired by scanning the entire field of view (512 lines/frame) at 15.49 Hz. After confirming a measurable stimulus-evoked response (>mean+(3*SD) of baseline), L2/3 glutamate transients were temporally resolved using limited area scans (100 lines/frame at 79.80 Hz).

### Data quantification and analysis

#### Whole-cell electrophysiology

All recordings were acquired at 20kHz and filtered at 2kHz (lowpass) using a Multiclamp 700B amplifier. Analog data were digitized using Digidata 1330 digitizer and Clampex 9 software (Axon Instruments). Access resistance was monitored throughout recordings (5mV pulses at 50Hz). Recordings with access resistance higher than 25MΩ or with > 20% change in the access resistance were discarded from the analysis. 70% series resistance compensation was applied to recorded currents in the voltage-clamp setting. Offline data processing was done with Clampfit 10 (Axon Instruments).

#### Stimulus-evoked glutamate transients

Image stacks were converted to maximum intensity projections and regions of interest (ROIs) were determined using a binary map of all pixels with a grayscale value > 2X standard deviation of maximum intensity projection (avoiding the slice anchor and stimulation electrode if these present within the FOV). The mean fluorescence intensities of the initial 50 frames of each trial served as F_0_. Fluorescence intensity traces were collected as the average intensity of all pixels within the ROI and normalized as ΔF/F_0_. Peak amplitude, the area under the curve (AUC), and full width at the half-maximum response (FWHM) of the glutamate transients were calculated using MATLAB (findpeaks function).

#### Analysis and statistics

Data analysis was performed using GraphPad Prism 8 (GraphPad Software, San Diego, California), MATLAB R2020a (Mathworks), Stata (StataCorp), and Microsoft Excel (Microsoft Corp). The normality of distributions was determined using the D’Agostino-Pearson K2 test and Shapiro-Wilk test. Outliers were identified based on data distribution (Grubbs’ test for parametric data and 1.5*interquartile range for non-parametric data). Average values for individual cells were compared across genotypes using a two-tailed, unpaired t-test (parametric data) or Mann-Whitney U test (non-parametric data). Paired comparisons were done using a two-tailed, paired t-test. Normalized (%) distributions of individual events were compared across genotypes using a two-sample Kolmogorov–Smirnov (KS) test. Data representing multiple time points within the same slice were compared across groups using two-way ANOVA (Friedman’s test for non-parametric data) or within groups using one-way ANOVA (Kruskal Wallis test for non-parametric data). For multiple comparisons, Bonferroni’s correction or Tukey’s test was used post-hoc for equal sample sizes. To compare the kinetics of input current-voltage (I/V) and action potential frequency-input current (F/I) relationships, linear regression was fitted across datasets, and slopes were compared between genotypes. To estimate the size and replenishment rate of the readily releasable pool (RRP), we fitted linear regressions to steady-state cumulative amplitudes of E/IPSCs. Y intercepts (RRP size) and slopes (RRP replenishment rates) of the linear regression fit were measured for grand means for genotypes or individual neurons.^27,28^ For pharmacology experiments, comparisons between cells before and after drug treatment within a genotype were done by paired t-test; comparisons between control and drug treatment groups across genotypes were done by two-way ANOVA with Tukey’s post hoc test. Statistical significance was set at *P* < 0.05, \**P* < 0.05, \*\**P* < 0.01, \*\*\**P* < 0.001, \*\*\*\**P* < 0.0001.

### Data availability

The authors confirm that the data supporting the findings of this study are available within the article and its supplementary material. Raw data were generated at The University of Utah. Raw and derived data supporting the findings of this study are available on request from the corresponding author.

## Results

### CK1δT44A mice have hyperexcitable sensory cortical circuits despite hyperpolarized membrane potentials *in vivo*

Hyperexcitable cortical sensory circuits can render the cortex susceptible to SD^12,13^ as well as confer tactile and thermal hypersensitivity^29,30^, similar to that observed previously in CK1δ_T44A_ mice.^15^ Under anesthesia or in quiet wakefulness, cortical networks can oscillate between depolarized ‘up states’ and quiescent ‘down states’.^31,32^ Up/down states are regulated by the balance of excitatory and inhibitory synaptic activity. As such, these states dynamically modulate cortical circuit gain, as neurons are more likely to fire action potentials (APs) during up states and less likely during down states.^32,33^ We recorded L2/3 pyramidal neuronal membrane potential (V_m_) oscillations during up and down states *in vivo*, to characterize network-driven excitability in WT and CK1δ_T44A_ mice (Figure 1A).

**Figure 1.**
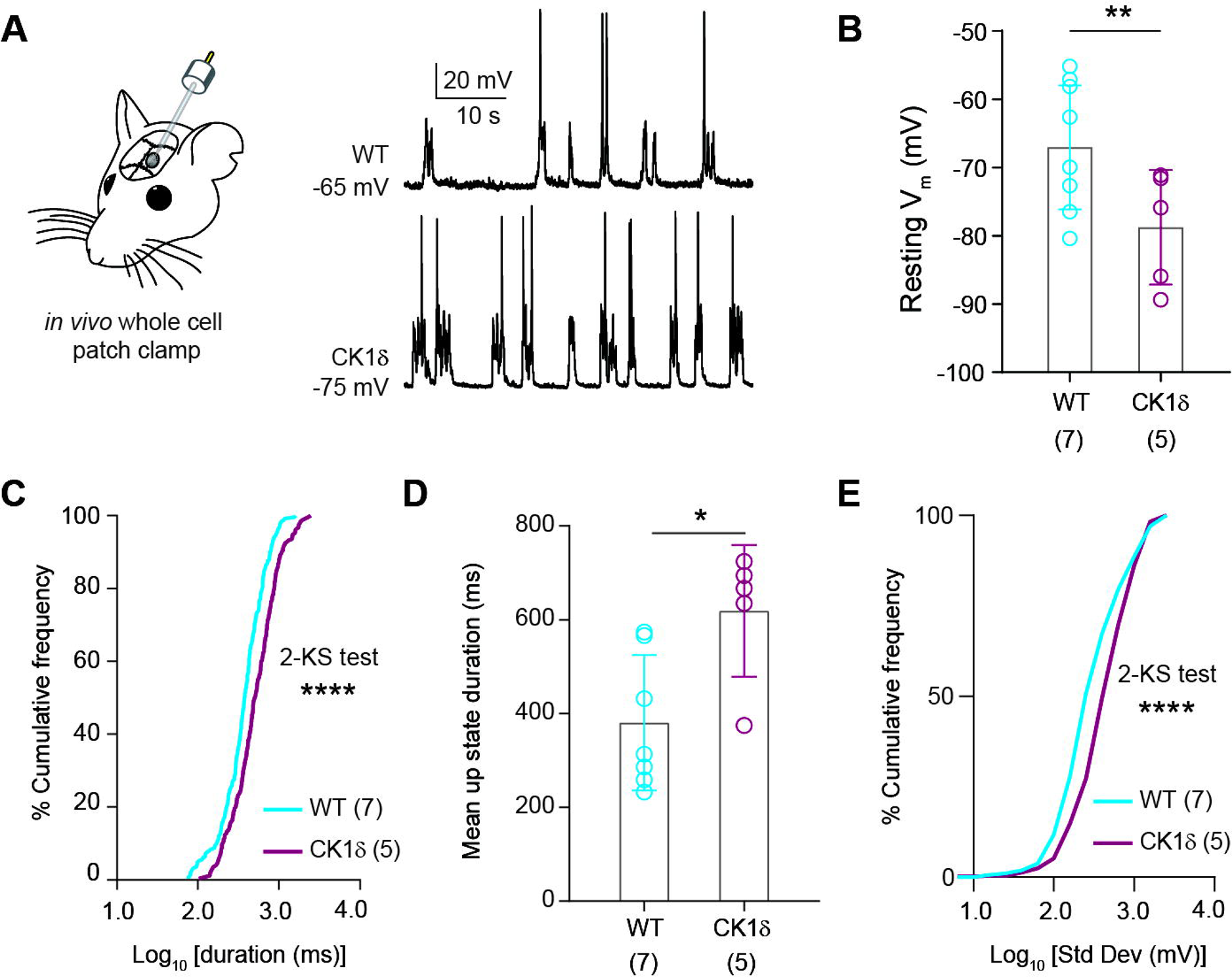
Increased up state duration and V_m_ variance found in CK1δ_T44A_ mice, despite hyperpolarized neuronal resting V_m_. (**A**) Schematic of *in vivo* whole-cell patch-clamp experiment along with representative current-clamp traces (L2/3 pyramidal neurons) of up and down states from WT and CK1δ_T44A_ mice. (**B**) CK1δ_T44A_ neurons were significantly hyperpolarized compared to WT. (**C**) Frequency histograms of upstate half-width show a positive shift in the upstate durations in CK1δ_T44A_ mice. (**D**) CK1δ_T44A_ mice show an increase in the mean up-state durations of individual animals and (**E**) an increase in membrane potential variance (standard deviation) during up states. Statistical analyses: unpaired t-test (B and D), 2-sample KS test (C and E,). Pooled data are means ± SD. *P < 0.05, **P < 0.01, and ****P < 0.0001. Exact P values can be found in table S1.

To our surprise, neuronal resting V_m_ during down states was significantly hyperpolarized (two-tailed t-test, *P<0.01*), in CK1δ_T44A_ mice – which is commonly associated with *reduced* excitability (Figure 1B). However, the duration of individual up states in CK1δ_T44A_ mice was significantly increased (2-sample KS test, *p<0.0001*, Figure 1C), as was the mean durations of all up states per animal (two-tailed t-test, *P<0.05*, Figure 1D). V_m_ variance, which represents the magnitude of persistent synaptic barrages during up states,^32^ was also significantly higher in CK1δ_T44A_ mice (2-sample KS test, *p<0.0001*, Figure 1E), confirming hyperexcitable cortical circuits in CK1δ_T44A_ mice, despite the hyperpolarized resting V_m_.

### Hyperpolarized membrane potential in CK1δT44A neurons is due to increased tonic inhibition

For further mechanistic dissection of the circuit phenotypes in CK1δ_T44A_ mice, we used *in vitro* whole-cell electrophysiology in acute cortical slices (Figure 2A). Consistent with our *in vivo* finding, CK1δ_T44A_ neurons were hyperpolarized compared to WT littermates (two-tailed t-test, *P<0.01*, Figure 2B), confirming the viability of brain slice preparation as a platform for more detailed examination.

**Figure 2.**
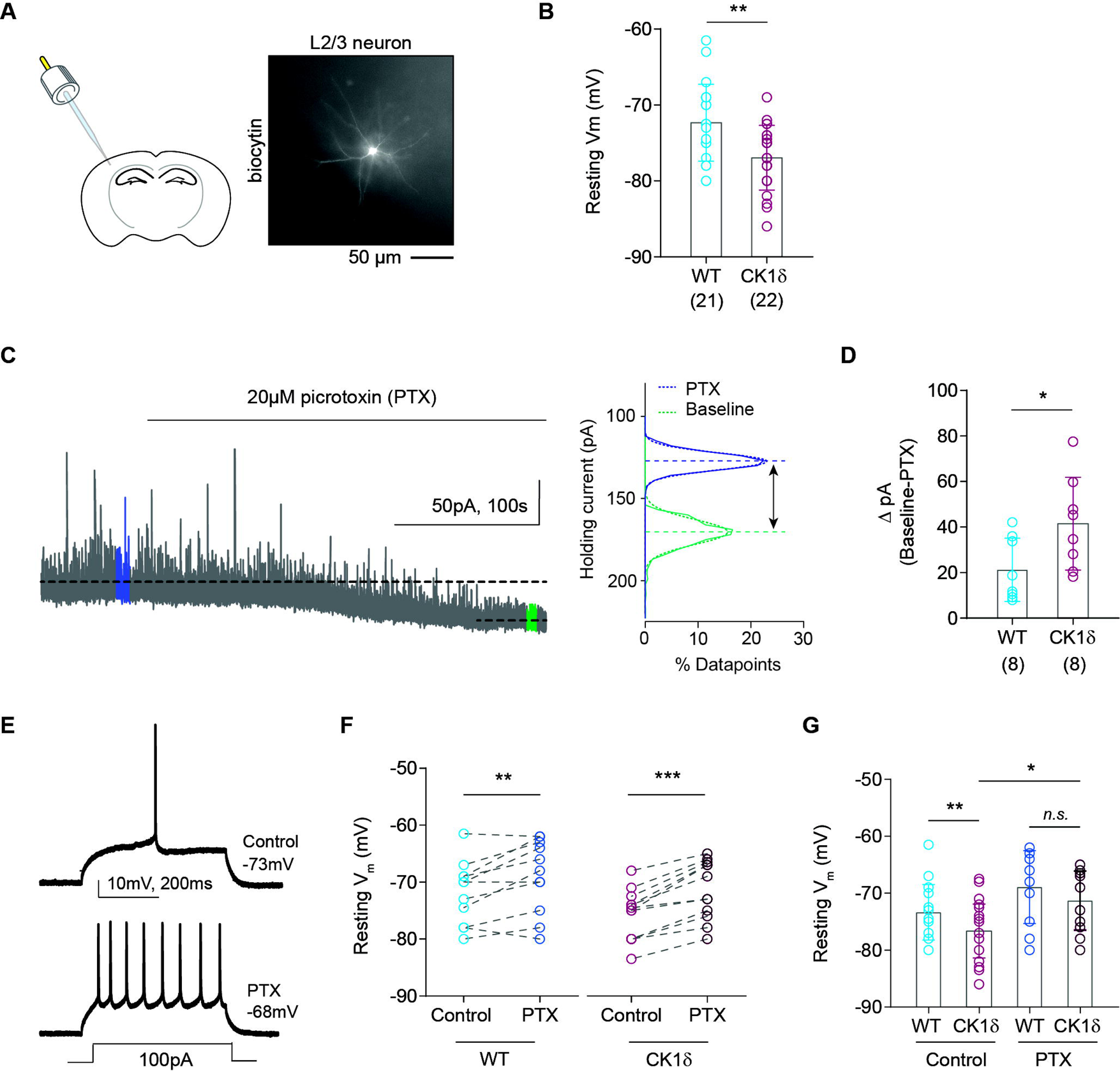
Increased tonic inhibition in CK1δ_T44A_ neurons primarily contributes to hyperpolarized resting V_m_. (**A**) Schematic showing *in vitro* whole-cell patch-clamp electrophysiology in acute coronal brain slices and a representative image of a typical biocytin-labeled excitatory L2/3 neuron. (**B**) Significantly hyperpolarized resting membrane potentials in CK1δ_T44A_ neurons compared to WT. (**C**) Representative trace from a WT neuron showing a reduction in tonic inhibitory current, after application of 20µM picrotoxin (GABA_A_ antagonist). Right: % distribution histograms along with gaussian fits (10^4^ datapoints) showing PTX sensitive holing current (tonic inhibitory current) as the difference between the % distribution means. (**D**) Tonic inhibitory currents were significantly larger in CK1δ_T44A_ neurons compared to WT. (**E**) Representative traces from a WT neuron showing resting membrane potential as well as voltage response to depolarizing current pulse before and after picrotoxin application. (F) The difference in the resting membrane potential between individual WT neurons as well as CK1δ_T44A_ neurons before and after picrotoxin treatment. (**G**) Pharmacological blockade of tonic inhibitory current lead to the ‘rescue’ of hyperpolarized membrane potentials in CK1δ_T44A_neurons. Statistical analyses: unpaired t-test (B), Mann Whitney test (D), paired t-test (F), and two-way ANOVA (G). Pooled data are means ± SEM. *P < 0.05, **P < 0.01, and ***P < 0.001. Exact P values can be found in table S1.

Tonic inhibitory currents can contribute to neuronal V_m_ hyperpolarization.^23^ We recorded tonic holding currents (holding potential; V_clamp_: 10mV) before and after blocking GABA_A_ receptors with 20 µM PTX (Figure 2C). We found that CK1δ_T44A_ neurons had significantly larger PTX-sensitive tonic inhibitory currents compared to WT (Mann Whitney test, *P<0.05*, Figure 2D). To determine if increased tonic inhibitory currents in CK1δ_T44A_ neurons substantially contribute to hyperpolarization, we recorded neuronal resting V_m_ before and after PTX treatment. As expected, PTX treatment depolarized neuronal resting V_m_ and reduced rheobase in both WT and CK1δ_T44A_ neurons (Figure 2E, S2). However, PTX-induced depolarization was significantly larger in CK1δ_T44A_ neurons compared to WT (Figure 2F). PTX also rescued neuronal V_m_ in CK1δ_T44A_ neurons (two-way ANOVA, Figure 2G), suggesting that increased tonic inhibition was the primary cause of resting V_m_ hyperpolarization.

### Increased frequency of evoked action potentials due to higher synaptic feedback excitation in CK1δT44A neurons

Thus far, our results show that CK1δ_T44A_ neurons were hyperpolarized due to increased tonic inhibition, yet there was an increase in cortical circuit excitability *in vivo*. Therefore, we examined whether CK1δ_T44A_ neurons were intrinsically hyperexcitable, by measuring membrane voltage (V_m_) responses to a range of input currents (Figure 3A). There was no difference in the neuronal membrane resistance (R_m_) measured below the action potential (AP) threshold (Figure S1), or the rheobase between the genotypes (Figure 3D). However, in response to larger input currents, CK1δ_T44A_ neurons fired APs at a significantly higher frequency compared to WT neurons (two-way ANOVA, Figure 3B), thus increasing the frequency-input current (FI) slopes in CK1δ_T44A_ neurons (Mann Whitney test, *P<0.05*, Figure 3C). Consistent with the FI slope data, the inter-spike interval between action potentials was significantly shorter in CK1δ_T44A_ neurons at larger input currents (two-way ANOVA, Figure 3H-J). Interestingly, there was no difference in AP half-width and after-hyperpolarization amplitude between the genotypes (Figure 3E-G), suggesting that altered AP kinetics did not contribute to the increased firing in CK1δ_T44A_ neurons. To determine a possible effect of increased tonic inhibition on suprathreshold activity in CK1δ_T44A_ neurons, we recorded evoked APs before and after PTX treatment (Figure 2E) and found no significant difference in FI slopes after PTX treatment in either genotype (Figure S2).

**Figure 3.**
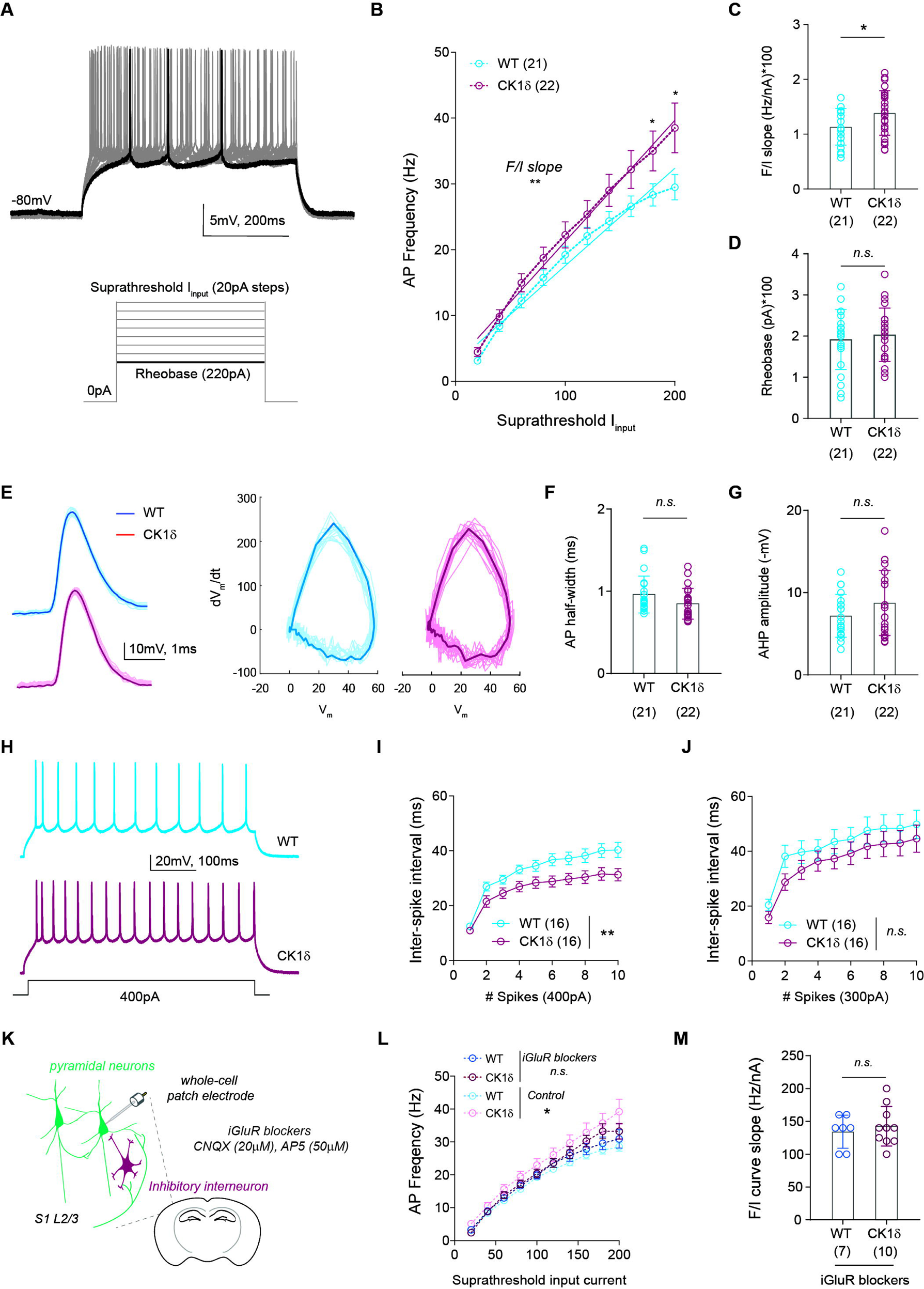
Increased evoked action potential frequency in CK1δ_T44A_ neurons was due to increased synaptic feedback excitation. (**A**) Representative traces of L2/3 excitatory neurons firing action potentials to suprathreshold current injections. (**B**) AP frequency in CK1δ_T44A_ neurons was significantly increased, especially at higher input currents along with the slopes of linear regression lines fitted to the FI plots. (**C**) Comparison of FI slopes quantified for individual neurons revealed that CK1δT44A had significantly higher FI slopes compared to WT. (**D**) No significant difference in rheobase between WT and CK1δT44A neurons. (**E**) Representative AP traces for WT and CK1δ_T44A_ neurons, as well as phase plots showing the rate of change of V_m_ during APs as a function of V_m_ (dV_m_/dt vs V_m_). (**F**) No difference was observed in AP half-width as well as (**G**) the after-hyperpolarization (AHP) amplitude. (**H**) Representative traces of WT and CK1δ_T44A_ neurons firing action potentials in response to 400pA current injection. (**I**) Comparison of inter-spike interval between WT and CK1δ_T44A_ neurons was significantly reduced following 400pA current injection, suggesting reduced spike frequency adaptation. (**J**) However, the inter-spike interval was not different in CK1δ_T44A_ neurons at 300pA. (**K**) Schematic showing pharmacological inhibition of postsynaptic glutamate receptors to block synaptic excitation during intracellular current injections. (**L**) Action potential frequency/input current (F/I) plots demonstrating that inhibition of postsynaptic glutamate receptors normalized the input-current-dependent increase in AP frequency seen in CK1δ_T44A_ neurons to WT levels. (**M**) No difference in the slopes of the linear regression fitted to the FI plots for individual neurons, between genotypes. Statistical analyses: Two-way ANOVA with Dunnett’s test for multiple comparisons (B, I and J), unpaired t-test (C, F, G, and M), ANCOVA (B), and Mann Whitney test (D). Pooled data are means ± SEM. *P < 0.05 and **P < 0.01. Exact P values can be found in table S1.

Neuronal action potential bursts elicited by >500ms long input currents, typically used in slice electrophysiology paradigms,^34,35^ can generate synaptic feedback excitatory currents within neocortical microcircuits.^36^ To test whether recurrent feedback excitation contributes to increased firing frequency in CK1δ_T44A_ neurons, we recorded evoked APs in the presence of ionotropic glutamate receptor antagonists (AMPARs and NMDARs) to abolish postsynaptic excitatory currents (Figure 3K). Glutamate receptor antagonists normalized the elevated AP frequency (two-way ANOVA, Figure 3L) and larger FI curve slopes in CK1δ_T44A_ neurons (two-tailed t-test, Figure 3M). Therefore, increased feedback synaptic excitation facilitated a higher frequency of evoked APs at larger input currents, suggesting that elevated excitatory synaptic activity may underlie the increased cortical excitability in CK1δ_T44A_ neurons.

### Baseline phasic excitatory and inhibitory neurotransmission is unaffected in CK1δT44A neurons

Balanced synaptic excitation and inhibition mediate “normal” cortical circuit excitability.^32,37^ We next examined whether baseline excitatory and inhibitory synaptic transmission contributed to hyperexcitability in CK1_δT44A_ circuits. We recorded AP-independent miniature excitatory and inhibitory postsynaptic currents (mE/IPSCs) in the presence of 1µM TTX (Figure S3A, S3B). We found that neither mEPSC amplitude nor frequency was significantly different between WT and CK1δ_T44A_ neurons (two-tailed t-test, Figure S3C, S3E). Similarly, mIPSC amplitude and frequency were not different between the genotypes (two-tailed t-test, Figure S3D, S3F). We found no difference in AP-dependent spontaneous excitatory and inhibitory neurotransmission (Figure S4). Finally, we recorded evoked E/IPSCs in L2/3 neurons in response to a single stimulus in layer 4 (L4). Again, we found no difference in the E/IPSC amplitude ratio between the genotypes (two-tailed t-test, Figure S4H).

NMDA receptor-mediated currents are a key component of excitatory neurotransmission and are modulated through phosphorylation by CK1 family proteins.^38^ To test if NMDA receptor-mediated currents were altered in CK1δ_T44A_ mice, we calculated the ratio of NMDA and AMPA components of responses (AMPA/NMDA ratio) evoked by a single stimulus (L4 to L2/3). We observed no significant difference in the AMPA/NMDA ratio in CK1δ_T44A_ compared to WT neurons (two-tailed t-test, Figure S4I). Taken together, our results suggest that neither phasic excitatory nor inhibitory synaptic transmission was altered in CK1δ_T44A_ neurons.

### Frequency-dependent adaptation deficit at CK1δT44A excitatory synapses

Thus far, we found that CK1δ_T44A_ mice exhibit an increase in local circuit excitability, yet there are no changes in phasic excitatory or inhibitory neurotransmission. We reasoned that this might be due to an altered postsynaptic response of CK1δ_T44A_ neurons to sustained synaptic stimulation.^32,39^ To test this, we recorded evoked E/IPSCs following short stimulus trains (L4-L2/3, 10 or 30 stimuli) at different frequencies (10, 20, 50Hz) (Figure 4A-B). To avoid polysynaptic responses, we verified the response latency and column-specificity of stimuli by confirming a lack of evoked responses outside the cortical column (Figure S7).

**Figure 4.**
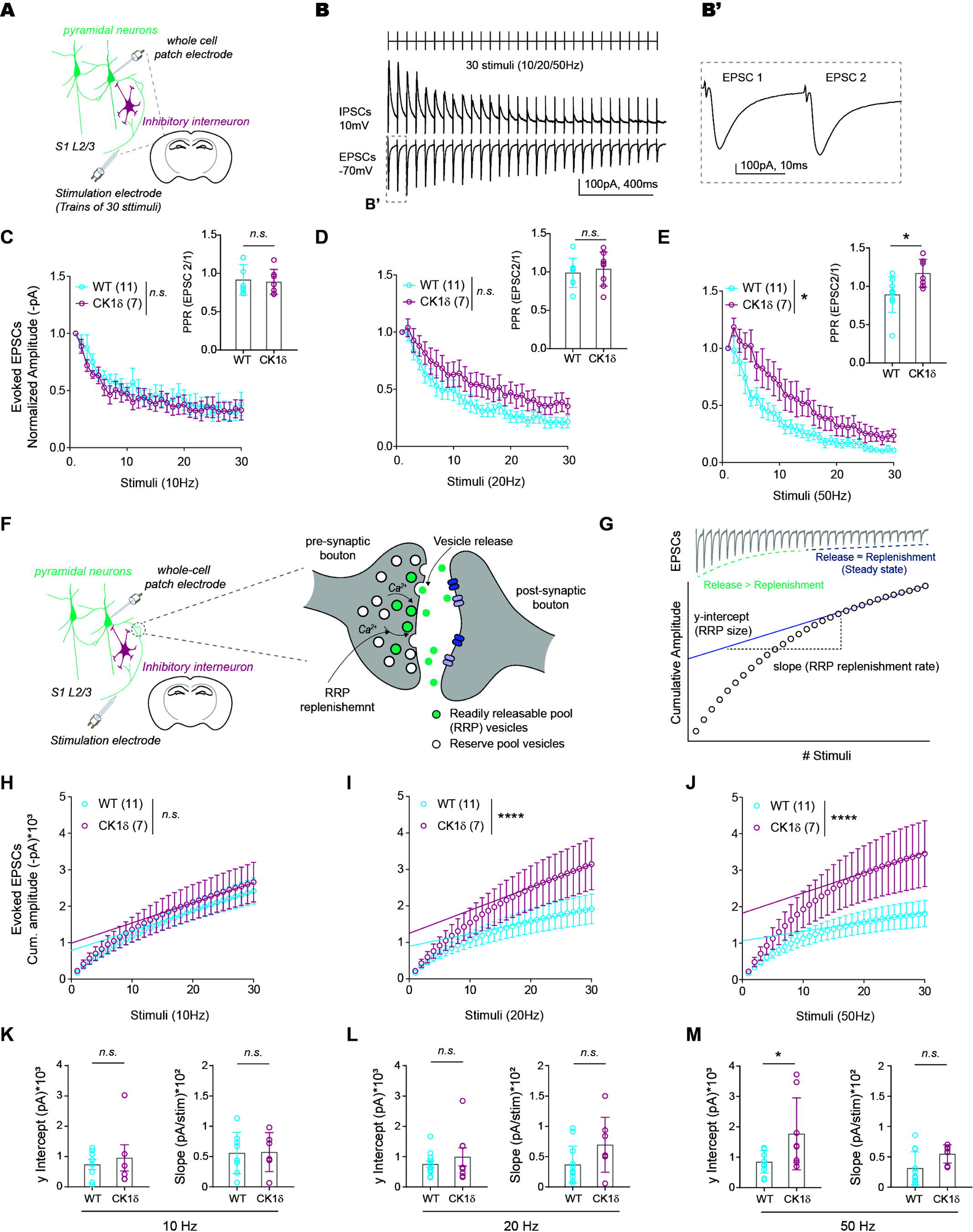
Stimulus-frequency-dependent adaptation deficits due to increased RRP size at CK1δ_T44A_ excitatory synapses. (**A**) Schematic representation of L2/3 cortical microcircuit showing the placement of stimulation as well as recording electrodes. (**B**) Representative traces showing evoked EPSC and IPSC responses to a train of 30 stimuli, at 20Hz. Traces of EPSCs evoked to the first two stimuli (**B’**), used for the quantification of paired-pulse ratio (PPR). (**C**) No significant difference between genotypes in the normalized EPSC amplitude as well as PPR at 10Hz and (**D**) 20Hz stimulus trains. (**E**) Normalized evoked EPSC response to 50Hz stimulus train was significantly higher at CK1δ_T44A_ synapse. *Inset:* significant difference in the paired-pulse ratio (PPR) at 50Hz between genotypes. (**F**) Schematic of a model synapse showing dynamic regulation of readily releasable pool (RRP) with simultaneous release and replenishment of synaptic vesicles upon repeated stimulation. (**G**) Schematic showing the analytical approach employed to explore the components of the presynaptic release machinery (RRP size and replenishment rate). (**H to J**) Cumulative amplitude plot for mean EPSC amplitudes: There was no significant difference in the slope of the linear regression fitted to the steady state between WT and CK1δT44A at any stimulus frequency. However, y-intercepts of the linear regression were significantly different between genotypes at both 20Hz (**I**) and 50Hz (**J**). (**K to M**) Slopes and y-intercepts for individual neurons: No significant difference in slopes, but significant difference in y-intercept at 50Hz (**M**). Statistical analyses: Repeated measures Two-way ANOVA with Dunnett’s test for multiple comparisons and unpaired t-test (C, D, and E), ANCOVA (H, I, and J), unpaired t-test or Mann Whitney u test (K, L, and M). Pooled data are means ± SEM. *P < 0.05 and ****P < 0.0001. Exact P values in table S1.

WT synapses showed the expected adaptation of postsynaptic responses to trains of 30 stimuli, as well as a paired-pulse ratio (PPR) consistent with that described in the literature for adult L4-L2/3 primary sensory cortex synapses.^40^ CK1δ_T44A_ neurons showed a similar response to WT at 10Hz stimulation. However, as stimulus frequency increased, the evoked EPSC adaptation showed a weak attenuation at 20Hz, a robust and significant reduction at 50Hz, (Two-way ANOVA, Figure 4C-E), and a significant increase in PPR (two-tailed t-test, Figure 4E, *inset*). We observed a similar frequency-dependent adaptation deficit in CK1δ_T44A_ EPSCs, following shorter duration (10 stimuli) trains (Figure S8). Interestingly, the impaired adaptation in CK1δ_T44A_ neurons was specific to excitatory currents, with evoked inhibitory currents exhibiting adaptation and PPR similar to WT at all stimulation frequencies (Two-way ANOVA, Figure S6A-C). Taken together, these results suggest a reduced adaptation of excitatory currents in CK1δ_T44A_ neurons following repeated high-frequency stimuli.

### Increased size of readily releasable pool impairs CK1δT44A excitatory synaptic adaptation

The rapid adaptation of excitatory postsynaptic currents is due to a presynaptic phenomenon involving Ca^2+^-mediated depletion and replenishment of the RRP of synaptic vesicles.^41^ The RRP size and its rate of replenishment influence the adaption of postsynaptic responses.^27,42,43^ We used the approach initially described by Schneggenburger *et al.*^44–46^ in the Calyx of Held, which was later adapted to cortical synapses by Abrahamsson *et al*.,^47^ to explore the components of the presynaptic release machinery that contribute to the reduced adaptation at CK1δ_T44A_ synapses. This model uses cumulative postsynaptic current amplitude as a function of stimulus number, with the slope of linear extrapolation of the steady-state indicating the RRP replenishment rate and the y-intercept corresponding to the RRP size (Figure 4F, G)^47^.

For EPSCs, there was no difference in the linear regression slope (replenishment rate) between genotypes at any of the frequencies tested. At 10Hz there was no difference in y-intercept (RRP size); however, at both 20Hz and 50Hz, there was a significant increase in the y-intercepts (more robust at 50Hz), consistent with a frequency-dependent increase in RRP size in CK1δ_T44A_ compared to WT animals (ANCOVA, *P<0.0001*, unpaired t-test, *p<0.05*, Figure 4H-M).

In contrast, neither replenishment rate nor RRP size was different for IPSC amplitudes between WT and CK1δ_T44A_ at any stimulus frequency (Figure S6D-I), confirming no difference in presynaptic adaptation at inhibitory synapses. These data predict that a larger presynaptic RRP during high-frequency stimulation at CK1δ_T44A_ glutamatergic synapses is responsible for the observed adaptation deficit.

### The adaptation deficit at CK1δT44A excitatory synapses is [Ca^2+^]e dependent

Recent evidence suggests that the RRP is a subset of fusion-competent and primed vesicles that may or may not be docked.^27^ Vesicle docking, priming, and release are Ca^2+^-dependent mechanisms, often modulated by extracellular [Ca^2+^],^44,47^ that ultimately determine the RRP size.^48,49^ Following high-frequency stimulation, residual Ca^2+^ and subsequent activation of biochemical cascades enhance vesicle priming and fusion, increasing the RRP size.^50^ We next tested whether increased RRP size at CK1δ_T44A_ glutamatergic synapses following high-frequency stimulation was dependent on [Ca^2+^]_e_. We evoked EPSC responses to stimulus trains (10, 20, and 50Hz) in the presence of low Ca^2+^ ACSF (0.65mM vs. 1.3mM in normal ACSF; Figure 5A, B). We found that the adaptation deficit at CK1δ_T44A_ excitatory synapses following 50Hz stimulation was ‘normalized’ with low Ca^2+^ ACSF (two-way ANOVA, Figure 5E), with RRP size (unpaired t-test, Figure 5I) and replenishment rate (unpaired t-test, Figure 5J) similar to WT levels. Therefore, the adaptation deficit at CK1δ_T44A_ excitatory synapses is dependent on [Ca^2+^]_e_, consistent with a presynaptic RRP-dependent mechanism.

**Figure 5.**
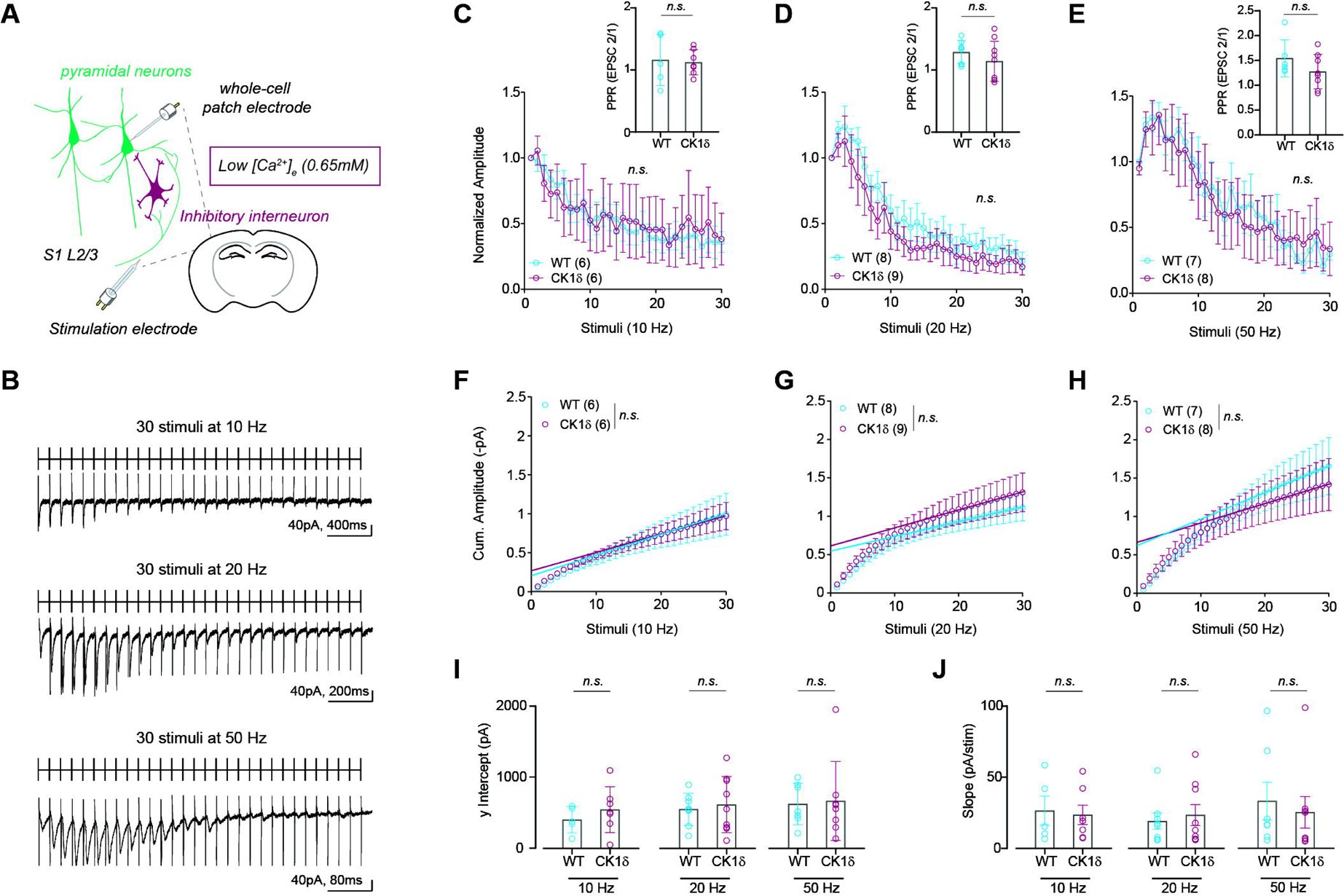
Impaired adaptation of evoked EPSCs at high stimulation frequencies was dependent on [Ca^2+^]_e_. (**A**) Schematic showing cortical microcircuits and experimental design to record evoked EPSCs in presence of low [Ca^2+^]_e_. (**B**) Representative race showing evoked EPSCs (at 10, 20, and 50Hz) recorded in low Ca^2+^ (0.65mM) ACSF, in response to a train of 30 stimuli. (**C to E**) In low [Ca^2+^]_e_ conditions, we found no difference in normalized EPSC amplitude and PPR between WT and CK1δ_T44A_ neurons at any frequency. (**F to H**) Similarly, we found no difference in the y-intercepts of linear regressions fitted to the steady-state cumulative EPSCs, in low [Ca^2+^]_e_ conditions between WT and CK1δ_T44A_ neurons. (**I and J**) In low [Ca^2+^]_e_ conditions, linear regression y-intercepts and the slopes of the cumulative EPSCs quantified for individual neurons were not different between genotypes for any stimulation frequency. Statistical analyses: repeated measures two-way ANOVA (C, D, and E), ANCOVA (F, G, and H), and unpaired t-test (C, D, E, I, and J). Pooled data are means ± SEM. Exact P values in table S1

### Larger glutamate transients following high-frequency stimulation in CK1δT44A slices

Our data thus far implicate that a larger RRP size at glutamatergic synapses during repeated high-frequency stimulation results in reduced adaptation. A larger RRP size would likely entail an increase in glutamate release, which can be measured with fluorescent indicators.^51^ Employing an approach similar to that described by Armbruster *et al*.,^26^ we imaged glutamate transients in L2/3 upon stimulation of L4 afferents, using two-photon laser-scanning microscopy (Figure 6A-B).

**Figure 6.**
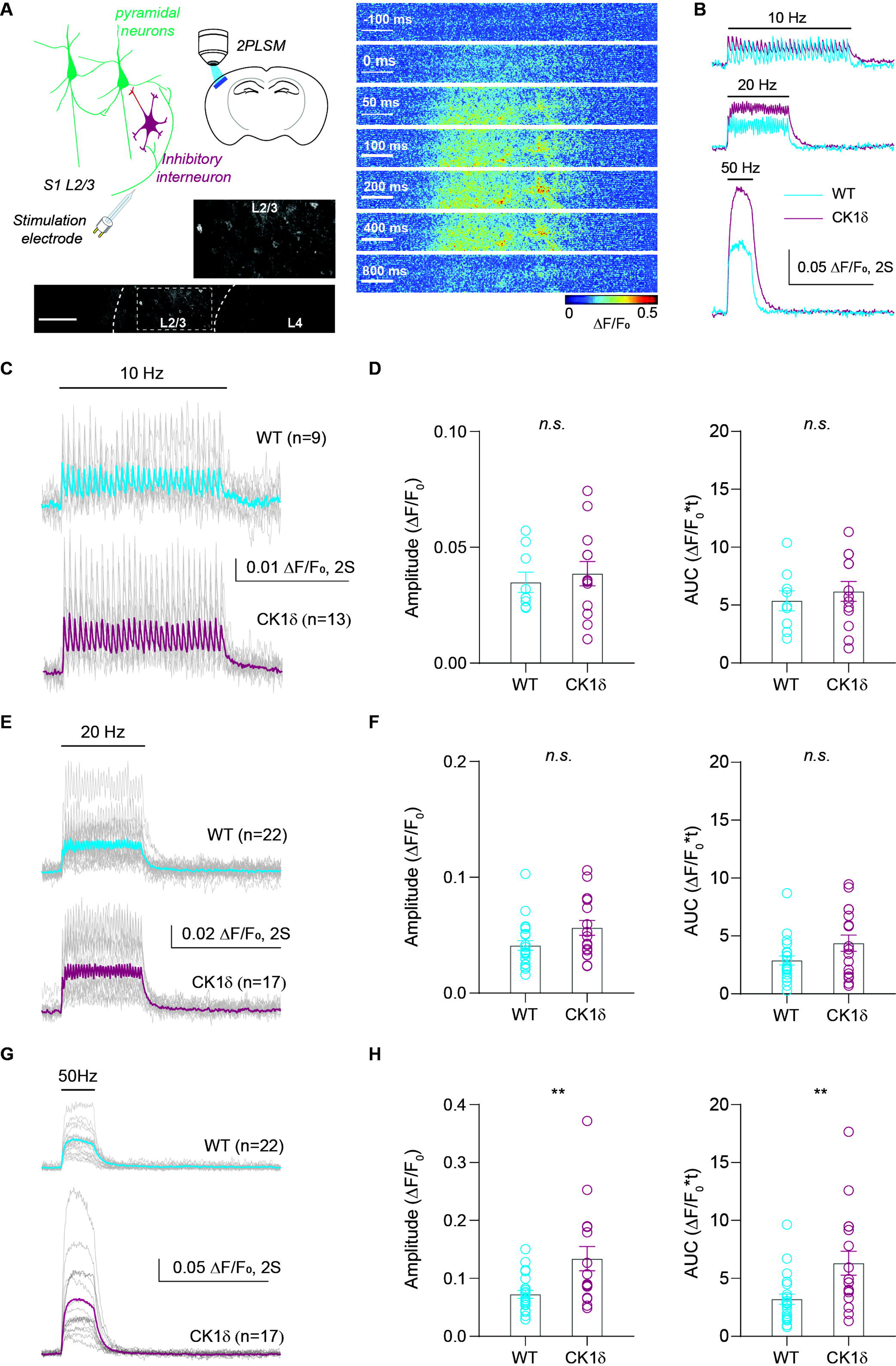
Larger glutamate transients in CK1δ_T44A_ slices evoked by high-frequency stimulation. (**A**) Schematic showing experimental design for two-photon fluorescent glutamate imaging in acute slices with a representative image of a limited area scan (100 lines/frame at 79.80 Hz) showing viral expression of iGluSnFR in neurons, (left). Image panels showing stimulus-evoked glutamate transient response over time, (50 Hz train of 30 stimuli, scale: 200µm, **Movie S2**). (**B**) Representative traces of evoked glutamate transient responses at 10, 20, and 50 Hz, in WT and CK1δ_T44A_ slices. (**C**) Traces of glutamate transient responses for individual slices (gray) and grand means (WT: cyan, CK1δ_T44A_: purple) evoked after 10Hz stimulation. (**D**) At 10Hz, the amplitude and area under the curve (AUC) of evoked glutamate transients were not significantly higher in CK1δ_T44A_ slices. (**E**) Traces of glutamate transient responses for individual slices and grand means evoked after 20Hz stimulation. (**F**) No difference in the amplitude or AUC of evoked glutamate transients evoked to 20Hz stimuli. (G) Traces of glutamate transient responses for individual slices and grand means evoked after 50Hz stimulation. (**H**) Both amplitude and AUC of glutamate transients evoked to 50Hz stimuli were significantly increased in CK1δ_T44A_ slices. Statistical analyses: Mann Whitney test (F and H) and unpaired t-test (D). Pooled data are means ± SEM. **P < 0.01. Exact P values in table S1.

Glutamate signals were evoked using a stimulation paradigm similar to that used for electrophysiology experiments. Glutamate maps acquired by scanning the entire field of view (at 15.49 Hz) showed a column-specific distribution of evoked responses, similar to evoked EPSC recordings (Figure S7, Movie S1). L2/3 glutamate transients in the appropriate column were then temporally resolved by faster limited-area scans (79.80 Hz at 100 lines/frame, Movie S2, *Materials and Methods*). Glutamate transients evoked in L2/3 in response to 10Hz and 20Hz stimulus trains were not significantly different between genotypes (Mann Whitney test, Figure 6C-F). However, there was a significant increase in peak amplitude and area under the curve (AUC) of glutamate transients in CK1δ_T44A_ slices compared to WT during 50Hz stimulations (Mann Whitney test, Figure 6G-H), suggesting increased glutamate release^52,53^ following high-frequency stimulation of CK1δ_T44A_cortical synapses. This finding offers convergent evidence of a presynaptic gain of function as a key underlying mechanism and a potential bridge between the cellular and circuit CK1δ_T44A_ phenotypes.

### An excitatory shift in the cortical circuits of CK1δT44A mice, *in vitro* and *in vivo*

Reduced adaptation at excitatory, but not inhibitory synapses, can result in a net excitatory shift in cortical microcircuits following stimulation, especially given the finding that reduced synaptic adaptation is associated with increased glutamate release. To test this hypothesis, we generated E/I ratios by measuring the amplitude of evoked E/IPSCs during stimulus trains (10 stimuli) recorded from the same neurons (Figure 7A). As expected, we observed a significant excitatory shift in CK1δ_T44A_ neurons after 50Hz stimulation (two-way ANOVA, *p<0.05*), but not at 20 or 10Hz (Figure 7B-D). We then tested whether the net excitatory shift upon intense synaptic stimulation is preserved in intact neural networks *in vi*vo (Figure 7E). During up states, cortical neurons exhibit both excitatory and inhibitory synaptic barrages when recorded using voltage-clamp electrophysiology *(Materials and Methods)*. We observed a robust increase in the duration of up-state evoked excitatory currents (Figure 7G), without a significant difference in the duration of inhibitory currents (Figure 7F), consistent with reduced adaptation observed exclusively at glutamatergic synapses in brain slices. Therefore, intense activity results in an excitatory shift in the output of cortical synapses *in vitro* and *in vivo,* in CK1δ_T44A_ mice.

**Figure 7.**
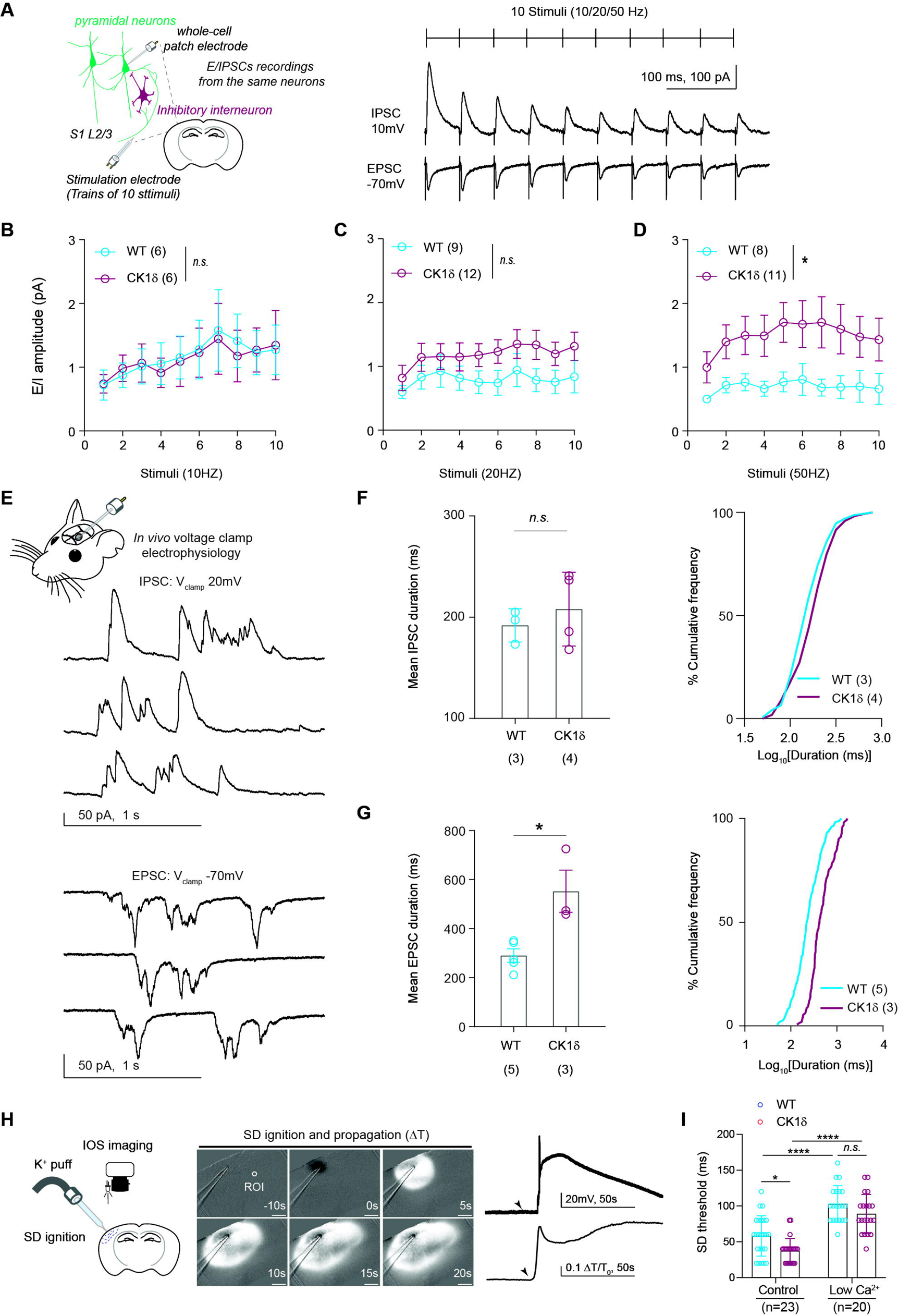
An excitatory shift in the cortical circuits found in CK1δ_T44A_ mice. (**A**) Schematic showing cortical microcircuit and experimental design along with representative traces of evoked E/IPSCs from a WT neuron in response to a train of 10 stimuli, recorded from the same neuron. (**B to D**) At 50Hz (**D**), E/IPSC amplitude ratios (E/I ratio) recorded from individual neurons show a significant shift towards excitation in CK1δ_T44A_ neurons. No difference in the E/I ratio between genotypes at 20Hz and 10Hz (**B** and **C**). (**E**) Representative traces showing inhibitory (V_clamp_ +20mV) and excitatory (V_clamp_ −70mV) synaptic barrages recorded in voltage clamp mode during an upstate *in vivo* (**F**) Comparison of IPSC half-width (animal means, left) shows no difference between WT and CK1δ_T44A_ mice. Frequency histograms of % distributions (right) show only a modest shift in half-width of the inhibitory events in CK1δ_T44A_ mice. (**G**) Comparison of EPSC half-width (animal means, left) shows a significant increase in CK1δ_T44A_ mice along with a frequency histogram of % distributions (right) showing a positive shift in half-width distribution of the excitatory events in CK1δ_T44A_ mice. (**H**) Schematic showing experimental design as well as an example image sequence showing the progression of a focally induced SD wave across space and time under control conditions (scale bar: 200 µm, **Movie S3**). Representative traces showing an increase in transmittance (ΔT/T_0_), obtained from an ROI 500µm from the induction site, as well as intracellular voltage rise recorded from different WT slices (arrows indicating the time of SD ignition). (**I**) CK1δ_T44A_ slices exhibit significantly reduced SD thresholds that were partially rescued in low [Ca^2+^]_e_ conditions. Statistical analyses: Repeated measures Two-way ANOVA (B, C, and D), Two-way ANOVA with Tukey’s test for multiple comparisons (I), Mann Whitney u test, and two-sample KS test (F and G). Pooled data are means ± SEM. *P < 0.05. Exact P values in table S1.

### Reduced [Ca^2+^]e normalizes SD susceptibility in CK1δT44A slices

CK1δ_T44A_ mice exhibit increased susceptibility to experimental SD induction *in vivo.*^15^ Similar observations of SD susceptibility in other migraine models are attributed to the increased release probability,^13^ or impaired clearance,^14^ of glutamate. To test whether the glutamatergic gain of function due to impaired adaptation at CK1δ_T44A_ excitatory synapses results in the facilitation of SD, we measured the threshold for SD induction *(Materials and Methods)* in acute cortical slices (Figure 7H, Movie S3). As expected, the threshold for SD initiation was significantly reduced in CK1δ_T44A_ slices compared to WT littermates (two-way ANOVA, Figure 7I).

To determine whether impaired adaptation at glutamatergic synapses had a causal role in the threshold difference, we measured SD thresholds in WT and CK1δ_T44A_ slices perfused with low Ca^2+^ ACSF, with the rationale that the reduced adaptation in CK1δ_T44A_ slices was [Ca^2+^]_e_ dependent (Figure 5). As SD induction itself depends on [Ca^2+^]_e_,^55^ low [Ca^2+^]_e_ significantly increased the SD threshold in slices from *both* genotypes, along with reduced maximum SD propagation (Figure S9). However, a comparison between genotypes in low [Ca^2+^]_e_ conditions revealed a ‘rescue’ of reduced SD thresholds in CK1δ_T44A_ slices (two-way ANOVA, Figure 7I) suggesting a [Ca^2+^]_e_ mediated mechanism underlies the reduced threshold in CK1δ_T44A_ compared to WT. Taken together, our results demonstrate that [Ca^2+^] _e_-mediated presynaptic adaptation deficit at excitatory synapses likely contributes to cortical network hyperexcitability and SD susceptibility in CK1δT44A mice (Figure 8).

**Figure 8.**
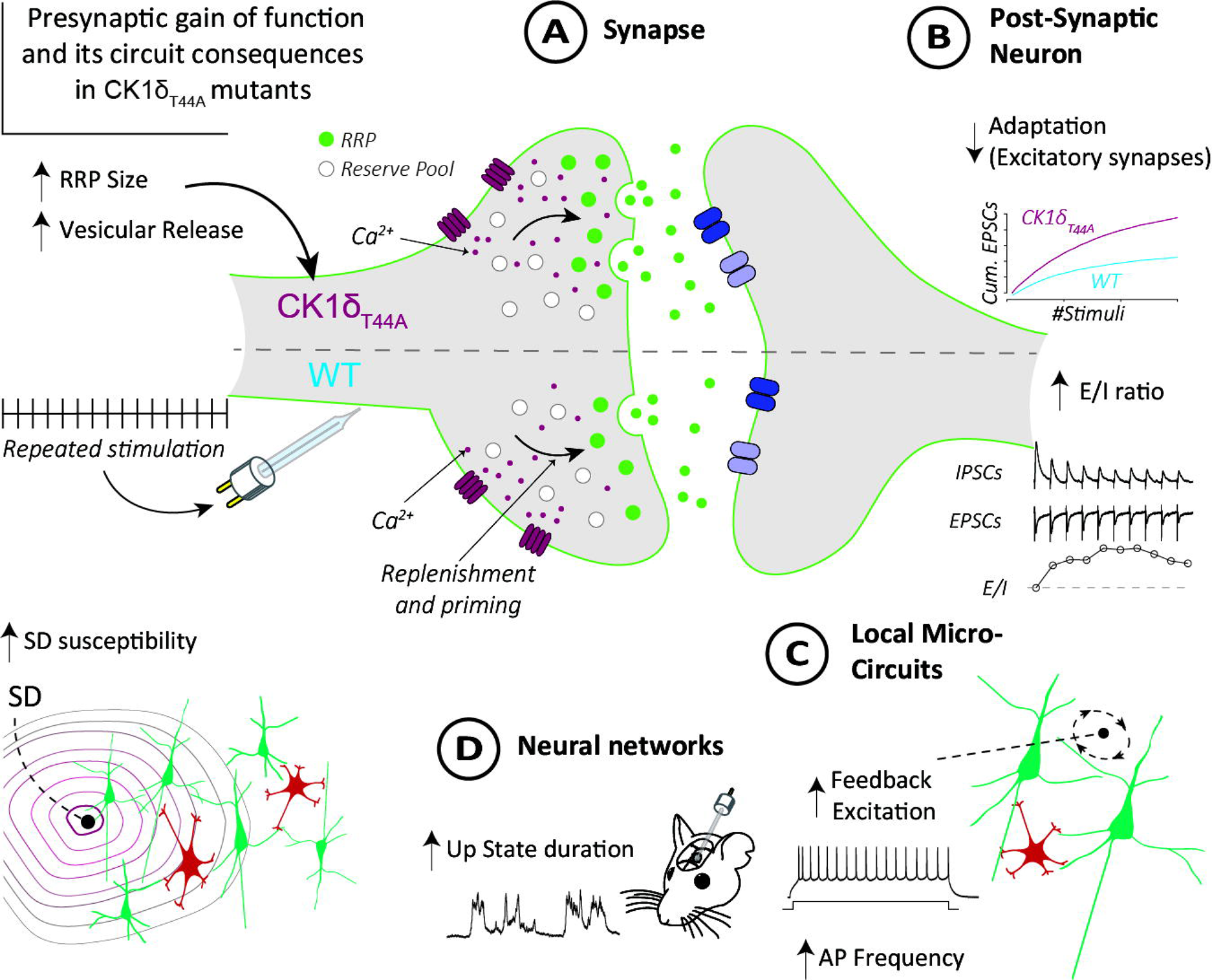
Model of cortical network excitability in CK1δ_T44A_ mice. **(A)** CK1δ_T44A_ excitatory (not inhibitory) synapses demonstrated reduced presynaptic adaptation to repeated high-frequency stimuli, mediated by a [Ca^2+^]_e_-dependent enhancement of RRP size, despite normal synaptic transmission at rest. **(B)** This led to an increase in the glutamate release and a resultant increase in cumulative amplitude of evoked EPSCs along with a higher excitation-to-inhibition ratio during sustained activity, both *in vivo* and in brain slices. **(C)** At the local microcircuit level, trains of action potentials elicited from single neurons increased glutamatergic feedback excitation in CK1δ_T44A_ compared to WT brain slices, further increasing firing frequencies at more intense input currents. **(D)** At a network level, CK1δ_T44A_ mice show an increase in the duration of up state activity *in vivo*, which is dependent on local circuit synaptic feedback excitation. Finally, SD susceptibility in CK1δ_T44A_ brain slices was returned to WT levels with the reduced [Ca^2+^]_e_, consistent with presynaptic contributions to the phenotype. Together, these findings show a presynaptic stimulus-dependent glutamatergic gain of function mediates a cortical network-level hyperexcitability phenotype in CK1δ_T44A_ mice.

## Discussion

Despite the pervasive and debilitating nature of migraine, the underlying mechanisms remain poorly understood.^1,2^ In CK1δ_T44A_ mice expressing a mutation found in humans with migraine with aura, we found a calcium-dependent reduction in presynaptic adaptation (Figure 4) and increased release (Figure 6) exclusively at glutamatergic synapses following intense circuit stimulation. This occurred despite increased tonic inhibition (Figure 2) as well as normal subthreshold and synaptic activity (Figure S1-S4). The stimulus-dependent increase in excitability bears an intriguing resemblance to migraine symptom descriptions and electrophysiological phenotypes in humans,^7,56,57^ and it provides a synaptic and circuit mechanism for the increased cortical excitability noted in CK1δ_T44A_ mice (Figure 1).^15^ Finally, we show a [Ca^2+^]_e_ -dependent increase in SD susceptibility CK1δ_T44A_ slices (Figure 7), directly implicating [Ca^2+^]_e_-dependent presynaptic mechanisms in migraine-relevant circuit excitability (Figure 8).^13,55^

Although migraine is more commonly polygenic,^3^ animal models of monogenic forms of the disorder offer a unique opportunity for mechanistic dissection.^12–14,58,59^ Recently, a loss of function mutation in the casein kinase 1 delta gene (CK1δ_T44A_) was identified in two families with a combination of familial migraine with aura and advanced sleep phase syndrome.^15,20^ Mice harboring this mutation showed migraine-relevant network excitability phenotypes including increased SD susceptibility and tactile and thermal hyperalgesia. These phenotypes were associated with increased c-fos expression in the trigeminal nucleus caudalis after administration of the migraine-triggering drug nitroglycerin.^15^ SD-associated vascular responses were also heightened in CK1δ_T44A_ mice. As SD likely triggers craniofacial pain at least partly through activation of vascular afferents,^9,60^ this phenotype might be evidence of a larger perturbation of pain-sensitive structures. The combined cortical excitability, vascular, and pain phenotypes in CK1δ_T44A_ mice,^15^ strongly argue for its broad disease relevance, despite the rarity of the mutation.

Sensory amplifications, during and between attacks, are a key feature of migraine.^61–63^ Sensory amplifications correlate with headache frequency and disease severity^64–66^ and are associated with poor clinical outcomes.^67^ Interestingly, migraineurs also report a lack of habituation to repeated sensory stimuli.^8,68–70^ Presynaptic adaptation is one of the principal mechanisms regulating sensory habituation and gain modulation in sensory circuits.^71,72^ In the cortex, a precise balance between synaptic excitation and inhibition is important for establishing the timing and signal-to-noise ratio necessary for signal processing.^73^ Stimulus-evoked and spontaneous activity in the sensory cortex involves a stereotypical pattern of excitation followed by inhibition.^74–76^ Even under normal conditions, synaptic excitation adapts more slowly than inhibition following repeated stimulation in the sensory cortex,^77,78^ generating a temporal window for the sensory signal integration.^24,79^ Reduced adaptation at excitatory – but *not* inhibitory - synapses upon high-frequency stimulation in CK1δ_T44A_ cortical neurons can thus significantly enhance signal transmission, resulting in sensory amplification.

Our findings are quite consistent with a scenario of enhanced cortical excitation due to the selective deficit in the adaptation of CK1δ_T44A_ excitatory neurons. The adaptation deficit is caused and accompanied by increased glutamate release. Increased glutamate release is a likely reason for the glutamate-dependent increase in feedback excitation we observed upon high-frequency stimulation (Figure 3). It also likely accounts for the increased up-state duration and membrane potential variance (Figure 1), which represents local synaptic barrages during depolarized up-states.

Interestingly, impaired adaptation is also present in other genetic models of migraine (FHM1)^80,81^ and it also occurs after SD,^82^ potentially providing a common presynaptic mechanism for the habituation failure to repeated stimuli reported in migraineurs.^7^ However, it is also important to note differences between the gain phenotypes in different models, as these may also have clinical relevance. The presynaptic gain of function phenotype in FHM1 differs from that of the CK1δ_T44A_ model, as a single stimulus is sufficient to elicit larger amplitude postsynaptic excitatory responses in FHM1.^13^ In the case of the CK1δ_T44A_ mouse model, repetitive high-frequency stimuli are necessary to elicit a glutamatergic adaptation deficit. Similarly, astrocytic transporter currents evoked by single stimuli, that represent the rate of glutamate clearance, are slower in FHM2 mice.^14^ This appears to correlate with the differences in the severity of disease between FHM syndromes (associated with hemiplegic or paralytic auras) and CK1δ_T44A_ mutation carriers, who report the non-hemiplegic auras which are also more common in the general migraine population.^3,11^ The response to repetitive rather than single stimulation is also consistent with patient reports of triggering migraines with high-intensity sensory inputs.^8,69^

Sensory perception is a multi-synaptic, multi-circuit process, and the mechanisms underlying altered perception extend well beyond the single-synapse level. Therefore, understanding migraine as a disorder of sensory gain requires an examination of circuit-wide mechanisms *in vivo*. Cortical slow oscillations are a well-characterized form of dynamic gain modulation in sensory circuits:^83^ depolarized ‘up states’ increase the likelihood of AP generation,^33,84^ and have been proposed as coincidence detectors for feedforward signal transmission^85^ during quiet wakefulness. Interestingly, increased up state frequency and duration have also been reported as a marker for network excitability in epilepsy models.^86,87^ Both migraine and epilepsy patients show alterations in high- and low-frequency cortical and thalamocortical oscillations, and the low-frequency oscillations are proposed as correlates of up states.^32,56,88^ Up state phenotypes thus provide a cortical circuit mechanism that can potentially bind the phenotypes observed in both humans and animal models of migraine.

Our *in vivo* investigations of cortical slow oscillations showed increased up-state duration and membrane voltage variance in CK1δ_T44A_ mice, despite hyperpolarized resting membrane potentials at baseline. During up-states, we found that CK1δ_T44A_ neurons received excitatory post-synaptic currents for a significantly longer duration than WT neurons, tipping the excitatory-inhibitory balance towards excitation. These *in vivo* findings bear an interesting resemblance to the increased frequency of action potentials evoked using intense stimuli *in vitro* in CK1δ_T44A_ neurons as both phenomena are dependent on glutamatergic input and rely on local feedback excitation. Since up states are *spontaneous* rather than evoked events that can occur during quiet wakefulness,^32^ CK1δ_T44A_ circuits may be hyperexcitable at baseline, without exposure to repetitive high-intensity stimuli.

At the synaptic level, our results showed a Ca^2+^-dependent increase in RRP size in CK1δ_T44A_ mice following high-frequency stimulation at excitatory synapses. The RRP is functionally defined as a subset of fusion-competent and primed vesicles in a presynaptic bouton that are more easily released than the remaining vesicle population.^27,48,49^ The RRP is thus dynamic, and its size changes as it is depleted and replenished simultaneously during repeated stimulation.^27^ This dynamic modulation of RRP size is dependent on [Ca^2+^] ^44,46^ as well as stimulus frequency.^89,90^ An increase in RRP size, as we observe in CK1δ_T44A_ on intense stimulation, has been shown to increase stimulus-evoked release probability,^89^ which can enhance Ca^2+^ -dependent glutamate release.^91^

The high-frequency stimulus-dependent increase we observed in the RRP of CK1δ_T44A_ synapses prompts a comparison with post-tetanic potentiation (PTP), which is attributed to elevated residual Ca^2+^, or changes to release machinery, including vesicle priming.^42,92^ Residual Ca^2+^ following high-frequency stimulation has been shown to activate biochemical targets like protein kinase C, Munc13, synapsin, and calmodulin/CaM kinase II, all of which regulate the size of the RRP.^93–95^ Post-tetanic potentiation is dependent on enhanced Ca^2+^ sensitivity of the vesicle fusion process, mediated by protein kinase C and Munc13.^93,94^ Moreover, genetic manipulations that impair RIM- and Munc13-mediated vesicle priming affect the RRP size.^27,96,97^ Interestingly, CK1 kinases co-localize with synaptic vesicular markers and phosphorylate a subset of synaptic vesicle-associated proteins (SV2). ^18^ Phosphorylation of SV2_A_ protein by CK1 family kinases controls the retrieval of synaptotagmin-1, a calcium sensor for the SNARE complex mediating the release of synaptic vesicles.^28,98^

Development-dependent morphological changes in synapses can also affect vesicle release probability. Recent evidence from CK1δ loss of function mutants in C. elegans suggests a potential role in axon maturation and stabilization.^99^ Multiple studies have demonstrated the variations in the presynaptic RRP size and release efficacy at different stages of axonal maturation.^100,101^ Therefore, a developmental role of the kinase in CK1δ_T44A_ mice could contribute to the presynaptic phenotypes observed in this study.^18^ Taken together, these findings support a possible role of CK1δ in the regulation of presynaptic function and suggest avenues for further investigation to uncover the molecular mechanisms responsible for the CK1δ_T44A_ adaptation phenotype. Moreover, the reduced thresholds for SD induction in CK1δ_T44A_ acute slices were normalized by low extracellular Ca^2+^ (Figure 7), elucidating a functional link between presynaptic facilitation and migraine-relevant circuit excitability.

Intriguingly, we observed an increase in the tonic inhibitory currents of CK1δ_T44A_ neurons, leading to a hyperpolarized resting V_m_, which is generally consistent with a *reduction* in excitability. At first approximation, this result appears to counter the other cellular and circuit phenotypes and suggests a possible compensatory mechanism. However, there is an equally tenable alternative explanation. Tonic inhibitory currents are mediated by extra and peri-synaptic GABA_A_ receptors^23^ and can be elicited by low levels of ambient GABA, due to their high GABA affinity.^22,102^ Tonic inhibitory currents are modulated by vesicular GABA release as well as reuptake, and therefore their activity correlates with the intensity of local network activity.^22,103^ Under baseline conditions, GABA transporters responsible for reuptake remain close to equilibrium (reversal) potential.^104^ As a result, even moderate circuit activity (brief neuronal depolarization or bursts of action potentials) is sufficient for GABA transporter reversal, resulting in both vesicular as well as non-vesicular GABA release,^105,106^ both of which can contribute to tonic inhibitory currents.^107^ Thus, a plausible explanation of the increased tonic inhibitory currents in CK1δ_T44A_ neurons is that they are a direct *consequence* of the hyperexcitable cortical circuits wrought by the presynaptic gain of function, rather than a compensatory response. It should also be considered that tonic inhibition may even *contribute* to network excitability, as the resulting increase in [Cl^−^]_i_ can render GABA excitatory, due to depolarized GABA reversal potential.^108–110^ That said, tonic inhibition as a more regulated compensatory response to circuit excitability (e.g. via reductions in GABA transporter expression)^111^ cannot be ruled out. Ultimately, despite the increased tonic inhibition, CK1δ_T44A_ cortical circuits remain hyperexcitable.

In conclusion, we uncovered evidence of a presynaptic gain of function at glutamatergic synapses in CK1δ_T44A_ mice, due to a calcium-dependent increase in RRP size that results in increased synaptic glutamate release. This gain of function caused a stimulus-dependent adaptation deficit of glutamatergic synaptic activity and was accompanied by enhanced network activity both *in vitro* and *in vivo,* that contributed to SD susceptibility. Taken together, our findings in CK1δ_T44A_ mice confirm presynaptic origins of cortical circuit excitability associated with phenotypically common form of migraine (Figure 8).

## Abbreviations

CK1δ: casein kinase-1δ
FHM: familial hemiplegic migraine
SD: spreading depolarization
S1 L2/3: primary somatosensory cortical layer 2/3
RRP: readily releasable pool
E/IPSCs: excitatory/inhibitory postsynaptic currents

## Acknowledgements

We thank Drs. Jeremy Theriot, Patrick Parker, and Kate Reinhart for providing technical direction; Loren L. Looger, and the Howard Hughes Medical Institute for providing iGluSnFR reagents to the research community.

## Supplementary material

Supplementary material is available at *Brain* online

