## Supplemental information for "Increased presynaptic excitability in a migraine with aura mutation"

### Supplementary Materials

#### Materials and Methods

##### Viral Injections

iGluSnFR expression was targeted to S1 L2/3 neurons using injections of adeno-associated virus (AAV1.hSyn.iGluSnFr.WPRE.SV40, Penn Vector Core catalog # AV-1-PV2723) 2 – 3 weeks before imaging[48]. Animals were anesthetized using isoflurane (~5% induction, <1.5% for surgery) and fixed in a stereotaxic frame (David Kopf Instruments, Models 962, 923B). Bupivacaine (30µl, subcutaneous) was administered as a local anesthetic and Rimadyl (5 mg/kg, subcutaneous) was used for postoperative analgesia, if necessary. Using standard sterile practices, bilateral injections were performed through small burr holes placed at coordinates relative to bregma (2 mm posterior, 3.2 mm lateral, 200 µm depth). AAV was injected (750-1000 nL, measured using a Hamilton Gastight syringe) through a pulled glass pipette (0.5 to 1MΩ resistance) connected to a sterile, Leur-lock injection syringe via polyethylene tubing. Post-surgery, mice were typically housed with 1 – 2 littermates, though occasionally they were singly housed.

##### SD induction thresholds

Acute coronal slices containing the barrel cortex prepared from WT and $CK1\delta_{T44A}$ mice (2 to 3 months) were perfused with ACSF at a flowing rate of 3 ml/min. Pressure-ejection pulses of 3M KCl (0.5 bar) of increasing duration (at 8 min intervals in 20 ms steps) were applied through a glass micropipette (R = 0.5-0.72MΩ, placed over layer1 and/or 2/3) onto the slice surface, using a pressure ejection system (MPPI-2, Applied Scientific Instruments), until an SD was elicited[13,21]. SD was detected by monitoring the associated change in intrinsic optical signal (IOS). SD propagation velocity was calculated as the rate of change of IOS, at a distance of 500 µm from the pressure-ejection pipette tip.

##### Recording AMPA/NMDA ratio

For AMPA/NMDA ratio measurements, a concentric bipolar metal stimulating electrode (FHC#30214, tip diameter: 125/25 µm, length: 75mm) connected to a stimulus isolator (World Precision Instruments) was placed on L4 and L5a (~400µm distance from recording site). Post-synaptic evoked AMPA and NMDA receptor-mediated currents were recorded by stimulating L4 and L5a (1ms, 1 to 10mV stimuli). Stimulation intensity providing ~50% of maximum response with no failure was selected for further experiments. Evoked AMPA currents were isolated at -70mV. Evoked NMDA-mediated currents were isolated at 40mV in the presence of DNQX (50 µM) and picrotoxin (20 µM).

#### Results

**Effect of picrotoxin (PTX) on intrinsic cellular properties.**

Pharmacological blockade of synaptic inhibition (tonic and phasic) by picrotoxin (PTX) application led to a significant reduction in the rheobase (minimum current required to elicit an action potential) in both WT and CK1$\delta_{T44A}$ neurons (Fig S2A). However, the comparison between genotypes revealed no significant difference in the rheobase (Fig S2B). Similarly, PTX application had no significant effect on input resistance (Fig S2C) as well as the FI relationship (Fig S2D) between WT and CK1$\delta_{T44A}$ neurons.

**Postsynaptic responses to trains of 10 stimuli**.

Similar to the longer trains of 30 stimuli, evoked EPSC and IPSC responses were recorded in response to trains of 10 stimuli, delivered at different frequencies (10, 20, and 50Hz). Similar to longer trains, normalized EPSC responses evoked with shorter 50Hz and 20Hz stimulus trains were significantly higher at CK1$\delta_{T44A}$ excitatory synapse, with no significant difference between genotypes following short 10Hz trains. Interestingly, we found no significant differences in the paired-pulse ratios (PPR) following short stimulus trains delivered at the aforementioned frequencies (Fig S7D and E). We also found no significant difference in normalized IPSC amplitude and PPR, between WT and CK1$\delta_{T44A}$ neurons, evoked at all stimulus frequencies (Fig S7F to H), corroborating the phenotype observed with longer stimulus trains.

#### Supplementary Figures:

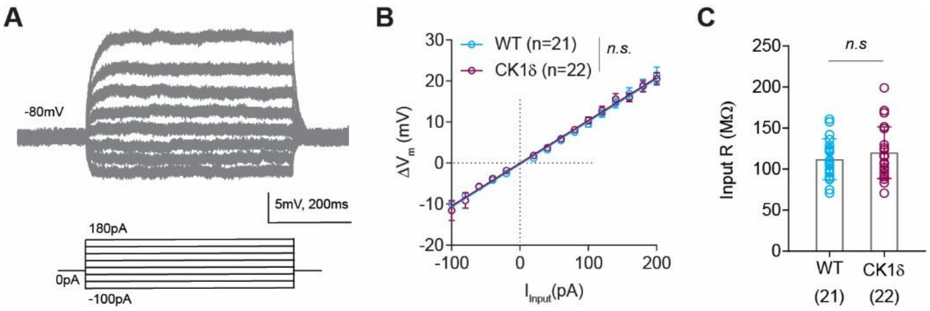


**Figure S1. Neuronal input resistance measurements.** (**A**) Representative traces showing membrane voltage responses to subthreshold current injections were recorded in current-clamp mode. (**B**) Slopes of the linear regression fitted to the IV curves of WT and CK1$\delta_{T44A}$ neurons were identical. (**C**) No difference between membrane resistance (IV-slopes) quantified for individual neurons, between WT and CK1$\delta_{T44A}$. Statistical analyses: ANCOVA (B) and unpaired t-test (C). Pooled data are means ± SEM. Exact P values can be found in Table S1.

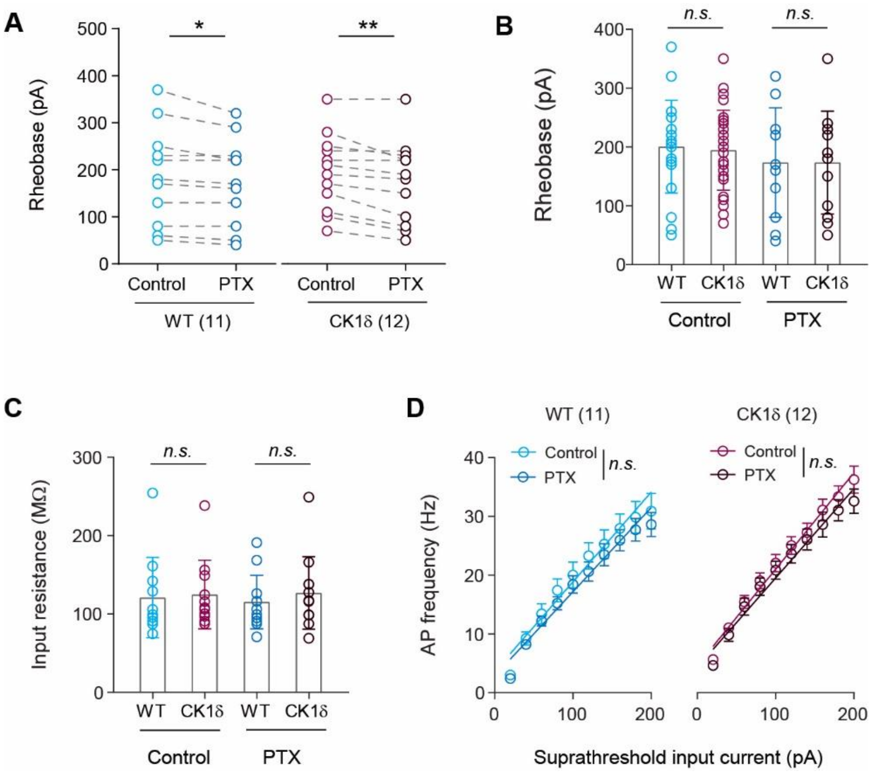


**Figure S2. Effect of picrotoxin (PTX) on intrinsic cellular properties**. (**A**) Pharmacological blockade of synaptic inhibition (tonic and phasic) by picrotoxin (PTX) application leads to reduced rheobase in both WT and CK1$\delta_{T44A}$ neurons (*p<0.01*, paired t-test). (**B**) However, PTX application did have a significant effect on rheobase *between* genotypes. (**C**) Similarly, PTX application did not have a significant effect on input resistance between WT and CK1$\delta_{T44A}$ neurons. (**D**) PTX application had no significant effect on action potential frequency upon suprathreshold current injections, in WT or CK1$\delta_{T44A}$ neurons. Statistical analyses: paired t-test (A), two-way ANOVA (B and C), and ANCOVA (D). Pooled data are means ± SEM. *$P < 0.05$, **$P < 0.01$. Exact P values can be found in Table S1.

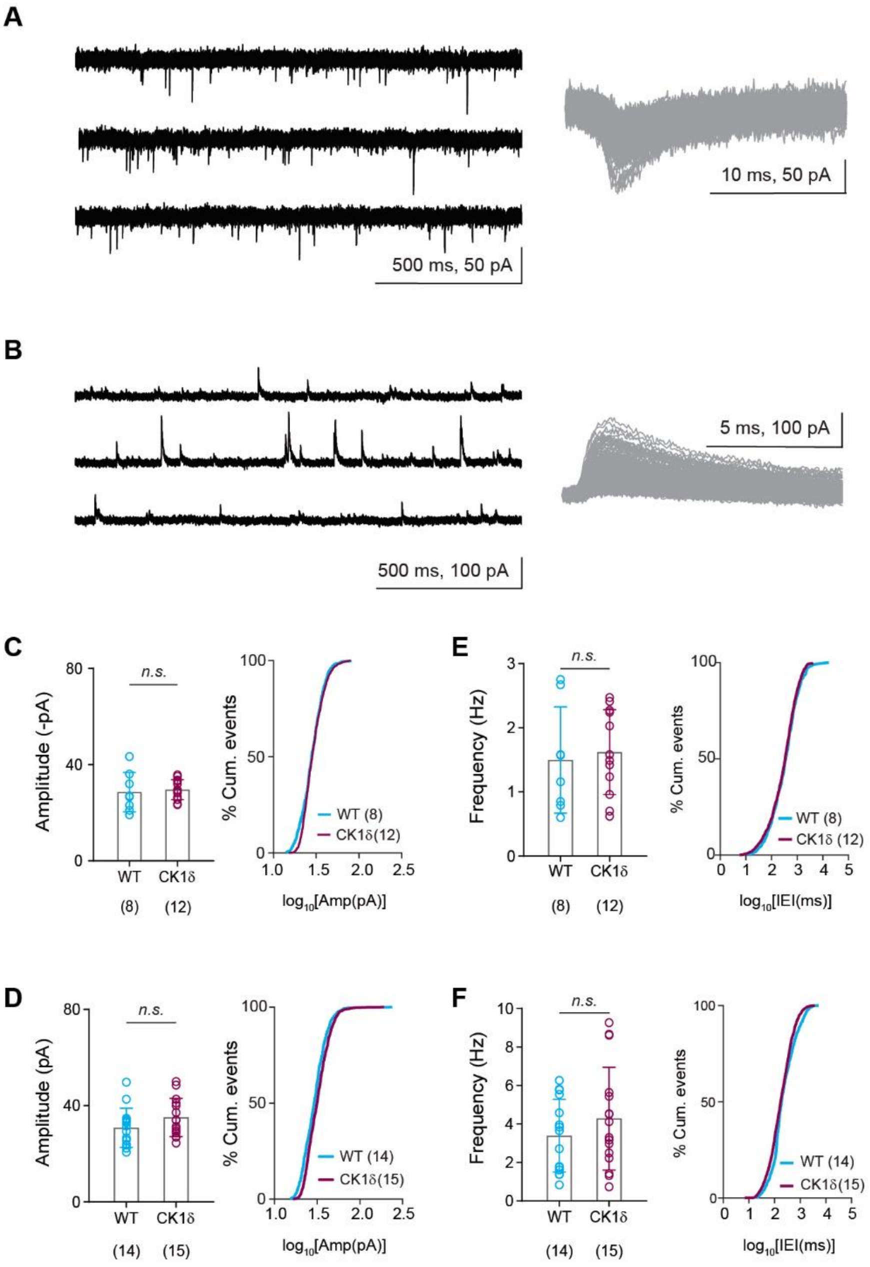


**Figure S3. Phasic synaptic neurotransmission is not altered in $CK1\delta_{T44A}$ neurons.** (**A**) Representative traces of miniature excitatory postsynaptic currents (mEPSC) as well as (**B**) miniature inhibitory postsynaptic currents (mIPSCs) in the presence of 1µM tetrodotoxin (TTX). The insets show traces of individual mEPSC and mIPSC events (gray) and respective means (black), recorded from a single neuron. No significant difference was found in the amplitude of mEPSCs (**C**) as well as mIPSC amplitudes (**D**) between $CK1\delta_{T44A}$ and WT neurons. (**E**) Comparison of mEPSC frequencies as well as (**F**) mIPSC frequency revealed no significant difference between WT and CK1$\delta_{T44A}$ neurons. Statistical analyses: unpaired t-test (C, D, E, F). Pooled data are means ± SEM. Exact P values can be found in Table S1.

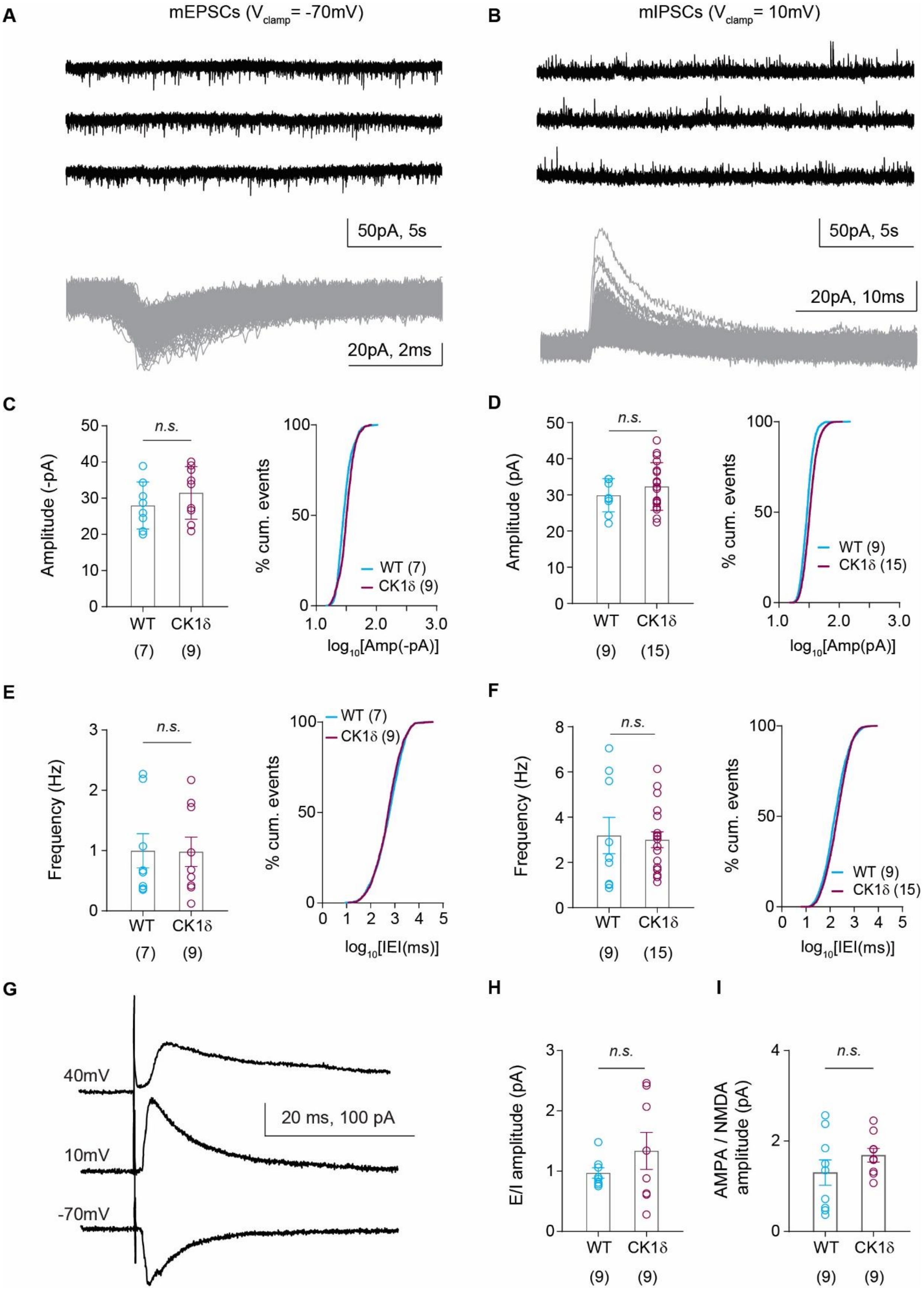
A
mEPSCs ($V_{clamp}$= -70mV)
50pA, 5s
20pA, 2ms
B
mIPSCs ($V_{clamp}$= 10mV)
50pA, 5s
20pA, 10ms
C
Amplitude (-pA)
n.s.
WT
CK1δ
(7)
(9)
% cum. events
WT (7)
CK1δ (9)
$\log_{10}$[Amp(-pA)]
D
Amplitude (pA)
n.s.
WT
CK1δ
(9)
(15)
% cum. events
WT (9)
CK1δ (15)
$\log_{10}$[Amp(pA)]
E
Frequency (Hz)
n.s.
WT
CK1δ
(7)
(9)
% cum. events
WT (7)
CK1δ (9)
$\log_{10}$[IEI(ms)]
F
Frequency (Hz)
n.s.
WT
CK1δ
(9)
(15)
% cum. events
WT (9)
CK1δ (15)
$\log_{10}$[IEI(ms)]
G
40mV
10mV
-70mV
20 ms, 100 pA
H
E/I amplitude (pA)
n.s.
WT
CK1δ
(9)
(9)
I
AMPA / NMDA amplitude (pA)
n.s.
WT
CK1δ
(9)
(9)

**Figure S4. Spontaneous E/IPSCs not altered in CK1$\delta_{T44A}$ neurons**. (**A**) Representative traces of spontaneous excitatory postsynaptic currents (sEPSC) as well as (**B**) spontaneous inhibitory postsynaptic currents (sIPSCs). The insets show traces of individual sEPSC and sIPSC events, recorded from a single neuron. (**C**) No significant difference was found in the amplitude of sEPSCs between CK1$\delta_{T44A}$ and WT neurons. (**D**) Similarly, sIPSC amplitudes were not significantly different between CK1$\delta_{T44A}$ and WT neurons. (**E**) Comparison of sEPSC as well as (**F**) sIPSC frequency, revealed no significant difference between WT and CK1$\delta_{T44A}$ neurons. (**G**) Representative traces of evoked AMPAR-mediated EPSC ($V_{clamp}$ -70mV), evoked $GABA_A$R-mediated IPSC ($V_{clamp}$ 10mV), and evoked NMDA current ($V_{clamp}$ 40mV) recorded from an S1-L2/3 excitatory neuron upon stimulation of L4/$L5_a$ afferents. (**H**) No difference was found either in the ratio of EPSC and IPSC amplitudes (E/I ratio) or (**I**) ratio of AMPAR and NMDAR mediated current amplitudes (AMPA/NMDA ratio) between WT and CK1$\delta_{T44A}$ neurons. Statistical analyses: unpaired t-test (C, D, E, F, H, and I). Pooled data are means ± SEM. Exact P values can be found in Table S1.

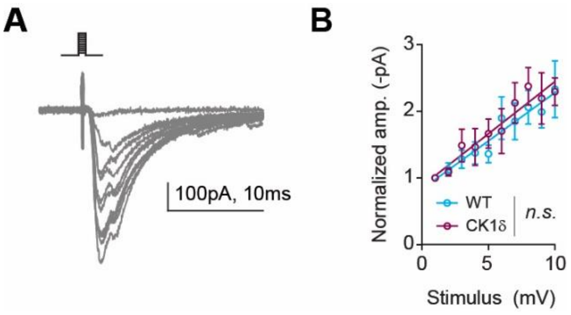


**Figure S5. No difference in the input-output function of evoked EPSCs between the genotypes.** (**A**) Representative traces evoked in response to single electrical stimuli with varying intensities (1 to 10mV), applied using a bipolar glass micropipette electrode. (**B**) The input-output relationship between WT and CK1$\delta_{T44A}$ to stimulation intensities used in this paradigm was not significantly different. Statistical analyses: ANCOVA (B). Pooled data are means ± SEM. Exact P values can be found in Table S1.

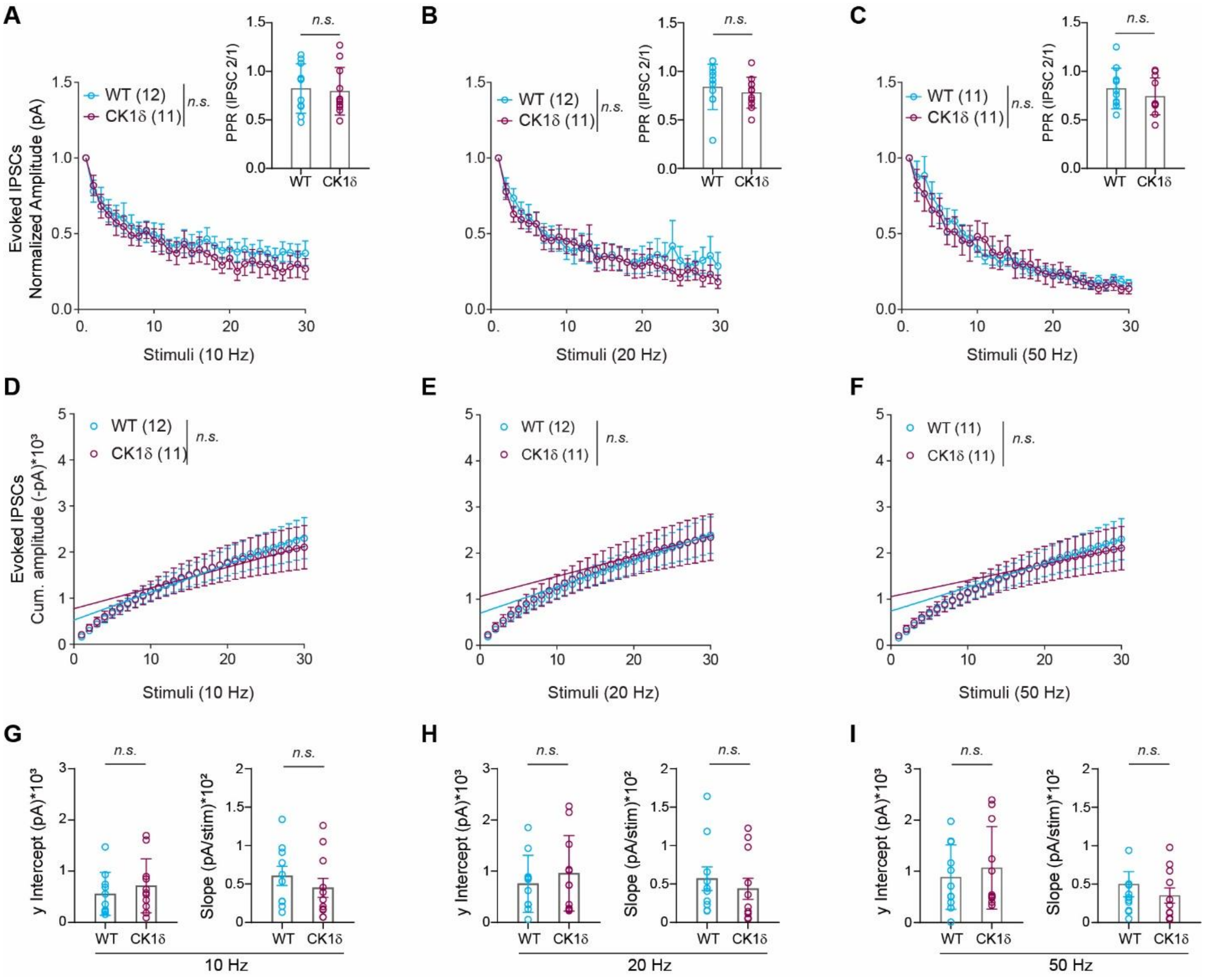


**Figure S6. Frequency-dependent adaptation deficit at CK1δ$_{T44A}$ excitatory synapses.** (**A to C**) No significant difference was found in normalized IPSC amplitude and PPR between WT and CK1δ$_{T44A}$ neurons. **(D to F)** The y-intercepts and the slopes of the linear regression fitted to the steady-state of mean cumulative amplitudes of evoked IPSCs were not different between genotypes for any stimulation frequency. (**G to I**) Similarly, linear regression y-intercepts and the slopes of the cumulative IPSCs quantified for individual neurons were not different between genotypes for any stimulation frequency. Statistical analyses: Two-way ANOVA with Dunnett's test for multiple comparisons and unpaired t-test (A to C), ANCOVA (D to F), unpaired t-test, or Mann Whitney u test (G to I). Pooled data are means ± SEM. Exact P values in table S1.

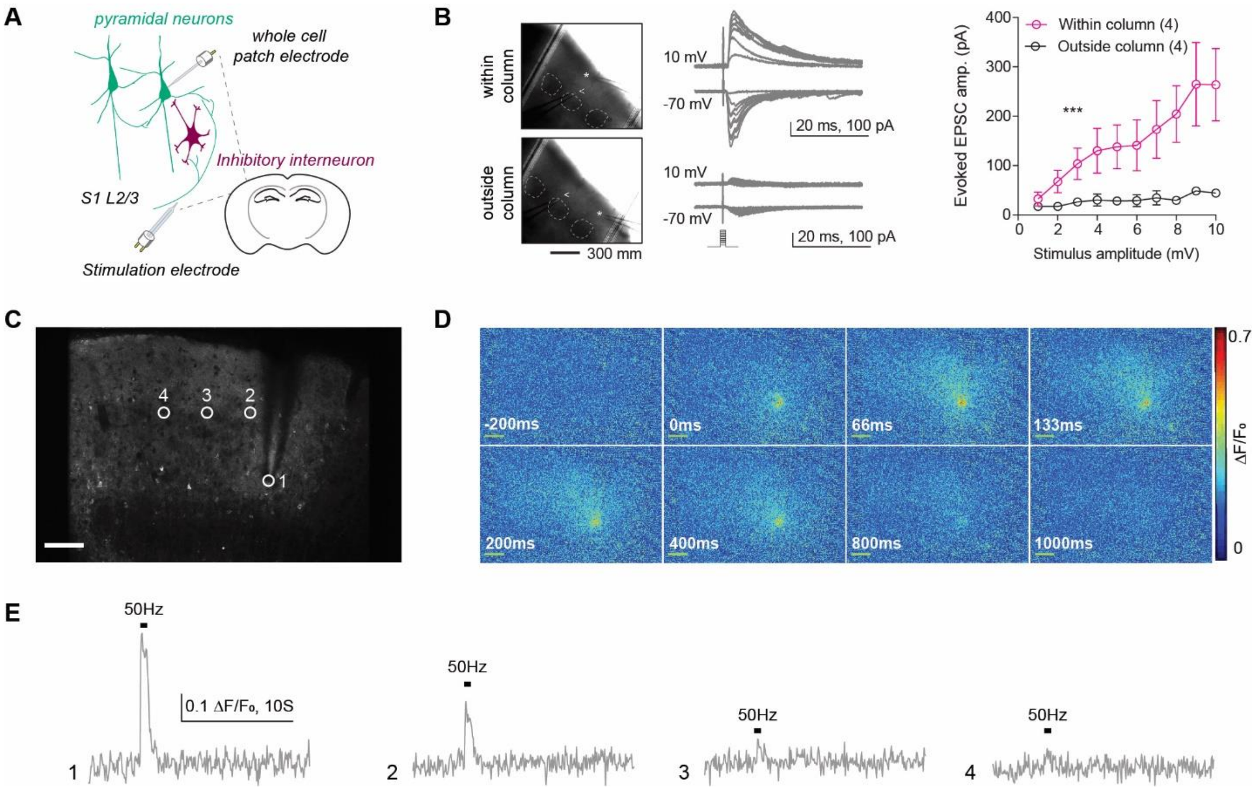


**Figure S7. Column specificity of the stimulation paradigm**. (**A**) Schematic showing cortical microcircuit and experimental setup for *in vitro* slice experiments. (**B**) *Left:* Images and representative traces showing experimentally evoked EPSC and IPSC responses recorded from neurons within (top) and outside (bottom) of a cortical column slice (scale bar: 200µm). *Right*: Quantification of EPSC amplitudes to increasing stimulus intensities for within-column and outside-column stimulation. (**C**) Representative image showing the placement of the stimulation electrode and ROIs across the slice (scale bar: 50µm). (**D**) Panels show stimulus-evoked (0ms, 50 Hz train of 30 stimuli) glutamate response over time, (change in fluorescence; $\Delta F/F_0$, **Movie S1**). (**E**) Traces of Glutamate response, captured from multiple ROIs over the slice, show diminished response away from the stimulation electrode. Statistical analyses: two-way ANOVA (B, *right*). Pooled data are means ± SEM. ***P < 0.001. Exact P values can be found in Table S1.

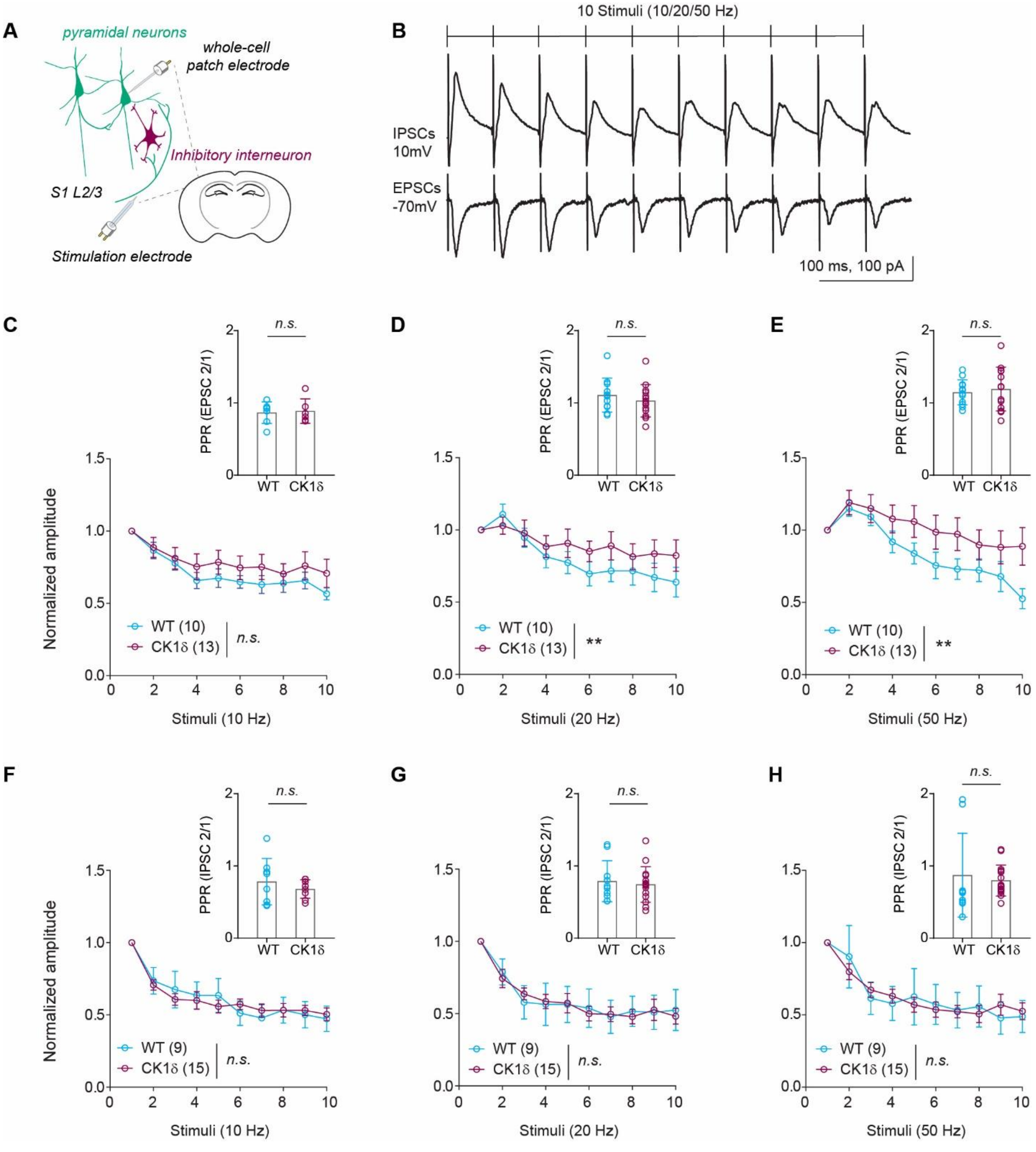


**Figure S8. Frequency-dependent adaptation at CK1δ$_{T44A}$ excitatory synapses following high-frequency trains of 10 stimuli.** (**A**) Schematic representation of L2/3 cortical microcircuit showing the placement of stimulation as well as recording electrodes. (**B**) Representative traces showing evoked EPSC and IPSC responses to a train of 10 stimuli, at 20Hz. Normalized evoked EPSC responses were significantly higher at CK1δ$_{T44A}$ synapse following to 50Hz and 20Hz stimulus trains. However, we found no significant difference in the paired-pulse ratio (PPR) at

50Hz between genotypes (**D** and **E**). No significant difference between the normalized EPSC amplitude as well as PPR following 10Hz (**C**) stimulus trains. (**F** to **H**) No significant difference was found in normalized IPSC amplitude and PPR between WT and CK1$\delta_{T44A}$ neurons. Statistical analyses: two-way ANOVA (C, D, E, F, G, and H) and unpaired t-test (insets: C, D, E, F, G, and H). Pooled data are means ± SEM. $^{**}P < 0.01$. Exact P values can be found in Table S1.

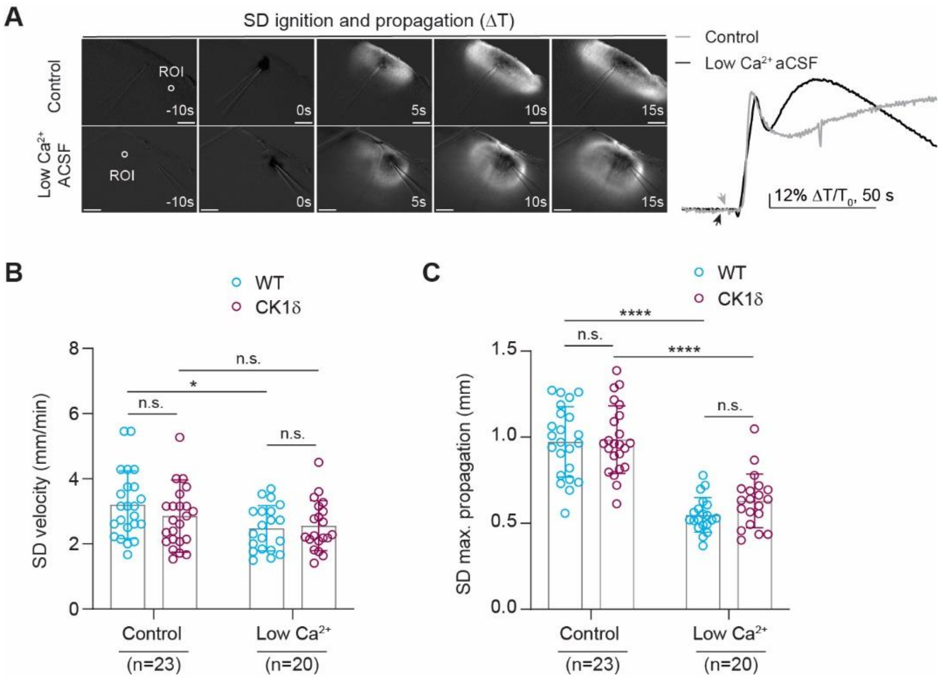


**Figure S9. Effects of low extracellular $Ca^{2+}$ on SD propagation.** (**A**) Example Image sequence (ΔT) showing the progression of a focally induced SD wave across space and time in control as well as low $[Ca^{2+}]_e$ conditions (scale bar: 200 μm), along with the representative traces of normalized transmittance signal (ΔT/T0), obtained from ROIs 500μm from induction site, in control and low $Ca^{2+}$ conditions (arrows indicating the time of SD ignition). (**B**) No difference in the SD propagation velocity between WT and $CK1\delta_{T44A}$ slices, under control conditions. Low $[Ca^{2+}]_e$ conditions resulted in a significant reduction in the SD velocity, only in WT slices. (**C**) No difference in the maximum SD propagation distance between WT and $CK1\delta_{T44A}$ slices, under control conditions. Low $[Ca^{2+}]_e$ conditions significantly reduced propagation distance, both in WT and $CK1\delta_{T44A}$ slices. Statistical analyses: Two-way ANOVA with Tukey's test for multiple comparisons (B and C). Pooled data are means ± SEM. *$P < 0.05$, ****$P<0.0001$. Exact P values can be found in Table S1.

#### Supplemental video files

**Movie S1. Stimulus evoked glutamate response related to Figure S6.** Example of a glutamate response evoked by a 50Hz stimulus, captured at a resolution of 918x596 µm (2x digital zoom) sampled at 15.49 Hz. The annotation (50Hz) indicates the onset and duration of the stimulus applied using a bipolar glass electrode. Left: raw image stack. Right: a pseudo color ΔF stack. The stack was cropped and gaussian smoothing (kernel: 3x3) was applied for clarity. Playback speed: 1× (15.49 Hz). Scale bar: 100 µm.

**Movie S2. Stimulus evoked glutamate response related to Figure 7.** Example of a glutamate response evoked by a 50Hz stimulus, captured using an area scan (100 lines/frame), at a resolution of 918x116.4 µm (2x digital zoom) sampled at 79.80 Hz. The annotation (50Hz) indicates the onset and duration of the stimulus applied through a bipolar glass electrode. Top: raw image stack. Bottom: a pseudo color ΔF stack. The stack was cropped and gaussian smoothing (kernel: 3x3) was applied for clarity. Playback speed: 1× (79.80 Hz). Scale bar: 100 µm.

**Movie S3. Example of SD induction threshold related to Figure 8.** A concatenated video demonstrating SD induction threshold in an acute slice perfused with regular ACSF, recorded as the IOS changes in response to high [$K^+$] puffs of increasing duration (20ms and 40ms). The raw images were acquired at a resolution of 1390x1040 µm, and sampled at 2 Hz. The annotations indicate the onset of the high [$K^+$] puff, applied through the pulled glass pipette. Left: a grayscale ΔT stack. Right: a pseudo color ΔT stack. The stack was cropped and gaussian smoothing (kernel: 3x3) was applied for clarity. Playback speed: 22.5× (45 Hz). Scale bar: 200 µm.

#### Table S1. Descriptive statistics.

| Figure | Test | Statistical value | P value | n | Definition of n |
|---|---|---|---|---|---|
| **Figure 1** | | | | | |
| 1B: Neuronal resting $V_m$ (*in vivo)* | Students t test, unpaired, two-tailed | t=2.371, df=12 | *p<0.05* | WT n=8, $CK1\delta_{T44A}$ n=5 | No. of mice |
| 1C: ‘up state’ duration: cumulative frequency distribution | 2-sample KS test | Kolmogorov-Smirnov D= 0.4652 | *p<0.0001* | WT n=8, $CK1\delta_{T44A}$ n=5 | No. of mice |
| 1D: Mean ‘up state’ duration | Mann Whitney test, two-tailed | U=3 | *p<0.05* | WT n=8, $CK1\delta_{T44A}$ n=5 | No. of mice |
| 1E: $V_m$ variance: cumulative frequency distribution | Students t test, unpaired, two-tailed | t=4.615, df=226 | *p<0.0001* | WT n=5, $CK1\delta_{T44A}$ n=5 | No. of mice |
| **Figure 2** | | | | | |
| 2B: Neuronal resting $V_m$ (*in vitro)* | Students t test, unpaired, two-tailed | t=3.241, df=41 | *p<0.01* | WT n=21, $CK1\delta_{T44A}$ n=22 | No. of neurons (1 neuron per acute slice) |
| 2D: tonic inhibitory current | Mann Whitney u | u=12.00 | *p<0.05* | WT n=8, $CK1\delta_{T44A}$ n=8 | No. of neurons (1 |

| | | | | | |
|---|---|---|---|---|---|
| | test, two-tailed | | | | neuron per acute slice) |
| 2F: effect of PTX on neuronal $V_m$ | Paired t test, two-tailed | t=3.301 df=10 | *p<0.01* | WT n=11 | No. of neurons (1 neuron per acute slice) |
| | Paired t-test, two-tailed | t=7.327 df=11 | *p<0.001* | CK1$\delta_{T44A}$ n=12 | No. of neurons (1 neuron per acute slice) |
| 2G: effect of PTX on neuronal $V_m$ | Kruskal-Wallis test, Dunn's multiple comparisons | Interactions<br>WT control vs CK1$\delta_{T44A}$ control, p=0.0041<br>WT PTX vs CK1$\delta_{T44A}$ PTX, p=0.3710<br>CK1$\delta_{T44A}$ PTX vs CK1$\delta_{T44A}$ control, p=0.0474 | *p<0.01* | PTX: WT=11, CK1$\delta_{T44A}$=12, Control: WT=21, CK1$\delta_{T44A}$=22 | No. of neurons (1 neuron per acute slice) |
| **Figure 3** | | | | | |
| 3B: F/I relationship | Two-way repeated measures ANOVA, Sidek's post-hoc test | Interactions:<br>Row p<0.0001 F = 331.9<br>Column p<0.0461 F = 4.218<br>Row x Column p<0.0001<br>F = 4.188 | *p<0.05* | WT n=21, CK1$\delta_{T44A}$ n=22 | No. of neurons (1 neuron per acute slice) |
| | ANCOVA | F (1, 393) =8.775 | *P<0.01* | | |

| | | | | | |
|---|---|---|---|---|---|
| 3C: rheobase | Students' t-test, unpaired, two-tailed | t=0.5363, df=41 | *p=0.59* | WT n=21, CK1$\delta_{T44A}$ n=22 | No. of neurons (1 neuron per acute slice) |
| 3D: F/I slope | Students' t-test, unpaired, two-tailed | t=2.290, df=45 | *p<0.05* | WT n=21, CK1$\delta_{T44A}$ n=22 | No. of neurons (1 neuron per acute slice) |
| 3F: AP half width | Mann Whitney u test, two-tailed | u=155 | *p=0.0.652* | WT n=21, CK1$\delta_{T44A}$ n=22 | No. of neurons (1 neuron per acute slice) |
| 3G: AP after-hyperpolarization | Students t test, unpaired, two-tailed | t=1.490, df=39 | *p=0.1442* | WT n=21, CK1$\delta_{T44A}$ n=22 | No. of neurons (1 neuron per acute slice) |
| 3I: inter-spike interval (400pA) | Two-way repeated measures ANOVA | Row F (1.142, 35.42) = 134.4 Column F (1, 31) = 6.023 Row x Column F (9, 279) = 3.021 | *p<0.01* | WT n=16, CK1$\delta_{T44A}$ n=16 | No. of neurons (1 neuron per acute slice) |
| 3J: inter-spike interval (300pA) | Two-way repeated measures ANOVA | Row F (1.062, 24.41) = 87.44 Column F (1, 23) = 1.041" Row x Column F (9, 207) = 0.6277" | *p=0.3182* | WT n=16, CK1$\delta_{T44A}$ n=16 | No. of neurons (1 neuron per acute slice) |
| 3L: F/I relationship | Two-way repeated | Interactions (GluR blockers) | *p=0.5347* | GluR blockers: | No. of neurons (1 |

| | | | | | |
|---|---|---|---|---|---|
| (GluR blockers) | measures ANOVA, Sidek's post-hoc test | Row *p<0.0001*, F=282.6 Column *p=0.5347*, F=0.4038 Row x Column *p=0.6960*, F=0.7131 | | WT=7, $CK1\delta_{T44A}$=10, Control: WT=21, $CK1\delta_{T44A}$=22 | neuron per acute slice) |
| 3M: F/I slope (GluR blockers) | Students t test, unpaired, two-tailed | t=0.5909, df=15 | *p= 0.5634* | WT n=7, $CK1\delta_{T44A}$ n=10 | No. of neurons (1 neuron per acute slice) |
| **Figure 4** | | | | | |
| 4C: Normalized EPSC amplitude (10Hz) | Two-way repeated measures ANOVA, | Stim x Genotype: F (29, 377) = 0.7501, *p=0.8244* Stim: F (29, 377) = 33.73, *p<0.0001* Genotype: F (1, 13) = 0.2266, *p=0.6419* | *p=0.8422* | WT n=11, $CK1\delta_{T44A}$ n=7 | No. of neurons (1 neuron per acute slice) |
| 4C: PPR (10Hz) | Students t test, unpaired, two-tailed | t=2.608, df=14 | *P=0.654* | WT n=11, $CK1\delta_{T44A}$ n=7 | No. of neurons (1 neuron per acute slice) |
| 4D: Normalized EPSC amplitude (20Hz) | Two-way repeated measures ANOVA, | Stim x Genotype: F (29, 406) = 0.6959. *p=0.8820* | *p=0.1391* | WT n=11, $CK1\delta_{T44A}$ n=7 | No. of neurons (1 neuron per acute slice) |

| | | | | | |
|---|---|---|---|---|---|
| | | Stim: F (3.879, 54.30) = 54.41, *p<0.0001* Genotype: F (1, 14) = 2.460, *p=0.1391* | | | |
| 4D: PPR (20Hz) | Students t test, unpaired, two-tailed | t=0.4819, df=14 | *p=0.3651* | WT n=11, CK1$\delta_{T44A}$ n=7 | No. of neurons (1 neuron per acute slice) |
| 4E: Normalized EPSC amplitude (50Hz) | Two-way repeated measures ANOVA, | Stim x Genotype: F (29, 435) = 1.626, *p=0.0227* Stim: F (3.096, 46.44) = 55.7, *p<0.0001* Genotype: F (1, 15) =5.538, *p=0.0327* | *p<0.05* | WT n=11, CK1$\delta_{T44A}$ n=7 | No. of neurons (1 neuron per acute slice) |
| 4E: PPR (50Hz) | Students t test, unpaired, two-tailed | t=2.608, df=14 | *p<0.05* | WT n=11, CK1$\delta_{T44A}$ n=7 | No. of neurons (1 neuron per acute slice) |
| 5I: Cumulative EPSC amplitude (20Hz) | two-way ANOVA | Stim x Genotype: F (29, 493) = 2.762, *p<0.0001* Stim: F (1.072, 18.22) = 38.54, *p<0.0001* | *p<0.0001* | WT n=11, CK1$\delta_{T44A}$ n=7 | No. of neurons (1 neuron per acute slice) |

| | | | | | |
|---|---|---|---|---|---|
| | | Genotype: F (1, 17) = 1.979, *p=0.1775* | | | |
| | ANCOVA (y-intercepts) | F (1, 201) = 26.25 | *p<0.0001* | | |
| 4H: Cumulative EPSC amplitude (10Hz) | two-way ANOVA | Stim x Genotype: F (29, 348) = 0.1001, *p>0.9999* Stim: F (1.164, 13.97) = 57.37, *p<0.0001* Genotype: F (1, 12) = 0.2280, *p=0.6416* | *p>0.9999* | WT n=11, CK1δ$_{T44A}$ n=7 | No. of neurons (1 neuron per acute slice) |
| | ANCOVA (y-intercepts) | F (1, 165) = 2.398 | *p=0.1234* | | |
| 4I: Cumulative EPSC amplitude (20Hz) | two-way ANOVA | Stim x Genotype: F (29, 493) = 2.762, *p<0.0001* Stim: F (1.072, 18.22) = 38.54, *p<0.0001* Genotype: F (1, 17) = 1.979, *p=0.1775* | *p<0.0001* | WT n=11, CK1δ$_{T44A}$ n=7 | No. of neurons (1 neuron per acute slice) |
| | ANCOVA (y-intercepts) | F (1, 201) = 26.25 | *p<0.0001* | | |
| 4J: Cumulative EPSC | two-way ANOVA | Stim x Genotype: F (29, 464) = 3.952, *p<0.0001* | *p<0.0001* | WT n=11, CK1δ$_{T44A}$ n=7 | No. of neurons (1 |

| | | | | | |
|---|---|---|---|---|---|
| amplitude (50Hz) | | Stim: F (1.061, 16.97) = 33.57, *p<0.0001* Genotype: F (1, 16) = 3.339, *p=0.0864* | | | neuron per acute slice) |
| | ANCOVA (y-intercepts) | F (1, 235) = 52.90 | *p<0.0001* | | |
| 4K: y-intercepts and slopes (10Hz) | Students t test, unpaired, two-tailed | t=0.5223, df=12 | *p=0.6110 (y-intercept)* | WT n=11, $CK1\delta_{T44A}$ n=7 | No. of neurons (1 neuron per acute slice) |
| | | t=0.07499, df=12 | *p=0.9415 (Slope)* | | |
| 4L: y-intercepts and slopes (20Hz) | Students t test (Mann Whitney's test), unpaired, two-tailed | t=0.8679, df=18 | *p=0.8065 (y-intercept)* | WT n=11, $CK1\delta_{T44A}$ n=7 | No. of neurons (1 neuron per acute slice) |
| | | u=24 | *p=0.1368 (Slope)* | | |
| 4M: y-intercepts and slopes (50Hz) | Students t test, unpaired, two-tailed | t=2.444, df=17 | *p<0.05(y-intercept)* | WT n=11, $CK1\delta_{T44A}$ n=7 | No. of neurons (1 neuron per acute slice) |
| | | t=2.085, df=16 | *p=0.0535 (Slope)* | | |
| **Figure 5** | | | | | |
| 5C: Normalized EPSC amplitude (10Hz) | two-way ANOVA | Stim x Genotype: F (29, 290) = 0.8232, *p=0.7292* Stim: F (2.161, 21.61) = 16.42, *p<0.0001* | *p=0.7292* | WT n=6, $CK1\delta_{T44A}$ n=6 | No. of neurons (1 neuron per acute slice) |

| | | | | | |
|---|---|---|---|---|---|
| | | Genotype: F (1, 10) = 0.005330, *p=0.9432* | | | |
| 5C: PPR (10Hz) | Students t test (Mann Whitney test), | t=0.2173, df=10 | *p=0.8234* | WT n=6, $CK1\delta_{T44A}$ n=6 | No. of neurons (1 neuron per acute slice) |
| 5D: Normalized EPSC amplitude (20Hz) | two-way ANOVA | Stim x Genotype: F (29, 435) = 0.3691, *p=0.9991* Stim: F (2.793, 41.90) = 36.66, *p<0.0001* Genotype: F (1, 15) = 0.7671, *p=0.3949* | *p=0.9991* | WT n=8, $CK1\delta_{T44A}$ n=9 | No. of neurons (1 neuron per acute slice) |
| 5D: PPR (20Hz) | Students t test (Mann Whitney test), | t=1.127, df=15 | *p=0.2774* | WT n=8, $CK1\delta_{T44A}$ n=9 | No. of neurons (1 neuron per acute slice) |
| 5E: Normalized EPSC amplitude (50Hz) | two-way ANOVA | Stim x Genotype: F (29, 377) = 0.4418, p=0.9952 Stim: F (3.352, 43.58) = 19.02, p<0.0001 Genotype: F (1, 13) = 0.01148 P=0.9163 | *p=0.9952* | WT n=7, $CK1\delta_{T44A}$ n=8 | No. of neurons (1 neuron per acute slice) |

| | | | | | |
|---|---|---|---|---|---|
| 5E: PPR (50Hz) | Mann Whitney u test, unpaired, two-tailed | u=14 | *p=0.2284* | WT n=7, CK1$\delta_{T44A}$ n=8 | No. of neurons (1 neuron per acute slice) |
| 5F: Cumulative EPSC amplitude (10Hz) | ANCOVA (y-intercept) | F (1, 177) = 0.004302 | *p=0.9484* | WT n=6, CK1$\delta_{T44A}$ n=6 | No. of neurons (1 neuron per acute slice) |
| 5G: Cumulative EPSC amplitude (20Hz) | ANCOVA (y-intercept) | F (1, 252) = 6.650 | *p=0.1005* | WT n=8, CK1$\delta_{T44A}$ n=9 | No. of neurons (1 neuron per acute slice) |
| 5H: Cumulative EPSC amplitude (50Hz) | ANCOVA (y-intercept) | F (1, 177) = 2.663 | *p=0.1045* | WT n=7, CK1$\delta_{T44A}$ n=8 | No. of neurons (1 neuron per acute slice) |
| 5I: y-intercept | Students t test (Mann Whitney test), unpaired, two-tailed | u=27 (50Hz), t=0.4248, df=15 (20Hz) t=0.8858, df=10 (10Hz) | *p=0.7789* (50 Hz), *p=0.6770* (20 Hz), *p=0.3965* (10 Hz) | WT n=8, CK1$\delta_{T44A}$ n=9 | No. of neurons (1 neuron per acute slice) |
| 5J: slopes | Students t test (Mann Whitney test), | u=24 (50Hz), u=35 (20Hz), t=0.2526, df=10 (10Hz) | *p=0.6943* (50 Hz), *p=0.9629* (20 Hz), | WT n=8, CK1$\delta_{T44A}$ n=9 | No. of neurons (1 neuron per acute slice) |

| | | | | | |
|---|---|---|---|---|---|
| | unpaired, two-tailed | | *p=0.8054* (10 Hz) | | |
| **Figure 6** | | | | | |
| 6D: 10Hz | Students t test, unpaired, two-tailed | t=0.5163, df=20 (amplitude) t=0.6369, df=20 (AUC) | *p=0.6113* (amplitude) *p=0.5314* (AUC) | WT n=9, CK1$\delta_{T44A}$ n=13 | No. of slices |
| 6F: 20Hz | Mann Whitney u test, two-tailed | u=121 (amplitude) u=139 (AUC) | *p=0.0631* (amplitude) *p=0.1806* (AUC) | WT n=22, CK1$\delta_{T44A}$ n=17 | No. of slices |
| 6H: 50Hz | Mann Whitney u test, two-tailed | u=86 (amplitude) u=89 (AUC) | *p<0.01* (amplitude) *p<0.01* (AUC) | WT n=22, CK1$\delta_{T44A}$ n=17 | No. of slices |
| **Figure 7** | | | | | |
| 7B: E/I amplitude (10Hz) | Two-way repeated measures ANOVA, | Stim x Genotype: F (9, 90) = 0.1893, *p=0.9948* Stim: F (1.169, 11.69) = 3.128, *p=0.0991* Genotype: F (1, 10) = 0.004637, *p=0.9471* | *p=0.9471* | WT n=6, CK1$\delta_{T44A}$ n=6 | No. of neurons (1 neuron per acute slice) |
| 7C: E/I amplitude (20Hz) | Two-way repeated measures ANOVA, | Stim x Genotype: F (9, 171) = 0.6830, *p=0.7236* Stim: F (9, 171) = 2.878, *p=0.0034* | *p=0.1628* | WT n=9, CK1$\delta_{T44A}$ n=12 | No. of neurons (1 neuron per acute slice) |

| | | | | | |
|---|---|---|---|---|---|
| | | Genotype: F (1, 19) = 2.108, *p=0.1628* | | | |
| 7D: E/I amplitude (50Hz) | Two-way repeated measures ANOVA, | Stim x Genotype: F (9, 144) = 1.486, *p=0.1583* Stim: F (1.181, 18.89) = 2.228, *p=0.1498* Genotype: F (1, 16) = 5.275, *p=0.0355* | *p<0.05* | WT n=8, CK1δ$_{T44A}$ n=11 | No. of neurons (1 neuron per acute slice) |
| 7F: EPSC half-width (animal means) | Mann Whitney u test, two-tailed | u=0 | *p<0.05* | WT n=5, CK1δ$_{T44A}$ n=3 | No. of mice |
| 7F: % distribution EPSC half-width | 2-sample KS test | Kolmogorov-Smirnov D= 0.4156 | *p=1.118e$^{-14}$* | WT n=5, CK1δ$_{T44A}$ n=3 | No. of mice |
| 7G: IPSC half-width (animal means) | Mann Whitney u test, two-tailed | u=8 | *p=0.7302* | WT n=5, CK1δ$_{T44A}$ n=4 | No. of mice |
| 7G: % distribution IPSC half-width | 2-sample KS test | Kolmogorov-Smirnov D= 0.2466 | *p= 6.050e$^{-05}$* | WT n=5, CK1δ$_{T44A}$ n=4 | No. of mice |
| 7I: SD threshold | Two-way ANOVA, Tukey's test | Interactions: | Genotype *p=0.0016,* Treatment | Control (WT n=24, | No. of slices |

| | | | | | |
|---|---|---|---|---|---|
| | for multiple comparison | WT control vs $CK1\delta_{T44A}$ control: *p=0.0248*<br>WT control vs WT low $Ca^{2+}$: *p<0.0001*<br>$CK1\delta_{T44A}$ control vs $CK1\delta_{T44A}$ low $Ca^{2+}$: *p<0.0001*<br>WT low $Ca^{2+}$ vs $CK1\delta_{T44A}$ low $Ca^{2+}$: *p=0.289* | *p<0.0001* | $CK1\delta_{T44A}$ n=23<br>Low $Ca^{2+}$ ACSF (WT, $CK1\delta_{T44A}$ n=20) | |
| **Supplemental Figures** | | | | | |
| **Figure S1** | | | | | |
| S1B: I/V relationship | ANCOVA (comparison of the slopes) | F=0.0002763<br>DFn=1<br>DFd=546 | *p=0.9867* | WT n=21, $CK1\delta_{T44A}$ n=22 | No. of neurons (1 neuron per acute slice) |
| S1C: input resistance | Students t test, unpaired, two-tailed | t=0.9182, df=40 | *p=0.3640* | WT n=21, $CK1\delta_{T44A}$ n=22 | No. of neurons (1 neuron per acute slice) |
| **Figure S2** | | | | | |
| S2A: rheobase | Paired t test, two-tailed | t=2.776 df=10 (WT)<br>t=3.952 df=11 ($CK1\delta_{T44A}$) | *p<0.05* (WT)<br>*p<0.01* ($CK1\delta_{T44A}$) | WT n=11, $CK1\delta_{T44A}$ n=12 | No. of neurons (1 neuron per acute slice) |
| S2B: rheobase | One-way ANOVA, Tukey's test | F= 0.4897<br>R squared= 0.02115 | *p=0.6906* | PTX: WT=11, $CK1\delta_{T44A}$=12, Control: | No. of neurons (1 neuron per acute slice) |

| | | | | | |
|---|---|---|---|---|---|
| | for multiple comparisons | | | WT=21, CK1$\delta_{T44A}$=22 | |
| S2C: input resistance | One-way ANOVA, Tukey's test for multiple comparisons | F=0.1559<br>R squared= 0.01101 | *p=0.9253* | PTX: WT=11, CK1δT44A=12, Control: WT=21, CK1δT44A=22 | No. of neurons (1 neuron per acute slice) |
| S2D: F/I relationship (slope) | ANCOVA | F=0.4464 (WT)<br>F=1.0432 (CK1$\delta_{T44A}$) | *p=0.5735* (WT)<br>*p=0.3084* (CK1$\delta_{T44A}$) | WT n=11, CK1$\delta_{T44A}$ n=12 | No. of neurons (1 neuron per acute slice) |
| **Figure S3** | | | | | |
| S3C: EPSC amplitude | Students t test, unpaired, two-tailed | t=0.3796, df=19 | *p=0.7084* | WT n=8, CK1$\delta_{T44A}$ n=12 | No. of neurons (1 neuron per acute slice) |
| S3D: IPSC amplitude | Students t test, unpaired, two-tailed | t=1.421, df=27 | *p=0.1667* | WT n=14, CK1$\delta_{T44A}$ n=15 | No. of neurons (1 neuron per acute slice) |
| S3E: EPSC frequency | Students t test, unpaired, two-tailed | t=0.3682, df=18 | *p=0.7170* | WT n=8, CK1$\delta_{T44A}$ n=12 | No. of neurons (1 neuron per acute slice) |
| S3F: IPSC frequency | Students t test, unpaired, two-tailed | t=1.059, df=29 | *p=0.2984* | WT n=14, CK1$\delta_{T44A}$ n=15 | No. of neurons (1 neuron per acute slice) |

| | | | | | |
|---|---|---|---|---|---|
| **Figure S4** | | | | | |
| S4C: mEPSC amplitude | Students t test, unpaired, two-tailed | t=1.036 df=15 | *p=0.3166* | WT n=7, CK1$\delta_{T44A}$ n=9 | No. of neurons (1 neuron per acute slice) |
| S4D: mIPSC amplitude | Students t test, unpaired, two-tailed | t=0.9865 df=25 | *p=0.3333* | WT n=9, CK1$\delta_{T44A}$ n=15 | No. of neurons (1 neuron per acute slice) |
| S4E: mEPSC frequency | Students t test, unpaired, two-tailed | t=0.04384 df=15 | *p=0.9656* | WT n=7, CK1$\delta_{T44A}$ n=9 | No. of neurons (1 neuron per acute slice) |
| S4F: mIPSC frequency | Students t test, unpaired, two-tailed | t=0.2538, df=25 | *p=0.8018* | WT n=9, CK1$\delta_{T44A}$ n=15 | No. of neurons (1 neuron per acute slice) |
| S4H: E/I ratio | Students t test, unpaired, two-tailed | t=1.144, df=14 | *p=0.2718* | WT n=8, CK1$\delta_{T44A}$ n=8 | No. of neurons (1 neuron per acute slice) |
| S4I: AMPA / NMDA ratio | Students t test, unpaired, two-tailed | t=1.207, df=16 | *p=0.2450* | WT n=8, CK1$\delta_{T44A}$ n=8 | No. of neurons (1 neuron per acute slice) |
| **Figure S5** | | | | | |
| S5B: Input/output relationship (linear | ANCOVA | F (1,195) = 0.09326 | *p=0.7604* | WT n=11, CK1$\delta_{T44A}$ n=7 | No. of neurons (1 neuron per acute slice) |

| | | | | | |
|---|---|---|---|---|---|
| regression slopes) | | | | | |
| **Figure S6** | | | | | |
| S6A: Normalized IPSC amplitude (10Hz) | Two-way repeated measures ANOVA, | Stim x Genotype: F (29, 609) = 0.5781, *p=0.9637* Stim: F (4.996, 104.9) = 40.15, *p<0.0001* Genotype: F (1, 21) = 0.4360, *p=0.5163* | *p=0.9637* | WT n=12, CK1$\delta_{T44A}$ n=11 | No. of neurons (1 neuron per acute slice) |
| S6A: PPR (10Hz) | Students t test, unpaired, two-tailed | t=0.2484, df=19 | *p=0.8065* | WT n=12, CK1$\delta_{T44A}$ n=11 | No. of neurons (1 neuron per acute slice) |
| S6B: Normalized IPSC amplitude (20Hz) | Two-way repeated measures ANOVA, | Stim x Genotype: F (29, 580) = 0.7966, *p=0.7684* Stim: F (2.865, 57.30) = 34.17, *p<0.0001* Genotype: F (1, 20) = 0.1341, *p=0.7181* | *p=0.7684* | WT n=12, CK1$\delta_{T44A}$ n=11 | No. of neurons (1 neuron per acute slice) |
| S6B: PPR (20Hz) | Students t test, unpaired, two-tailed | t=0.6841, df=19 | *p=0.5022* | WT n=12, CK1$\delta_{T44A}$ n=11 | No. of neurons (1 neuron per acute slice) |

| | | | | | |
|---|---|---|---|---|---|
| S6C: Normalized IPSC amplitude (50Hz) | Two-way repeated measures ANOVA, | Stim x Genotype: F (29, 580) = 0.520, *p=0.9829* Stim: F (3.328, 66.57) = 43.55, *p<0.0001* Genotype: F (1, 20) = 0.02994, *p=0.8644* | *p=0.9829* | WT n=12, $CK1\delta_{T44A}$ n=11 | No. of neurons (1 neuron per acute slice) |
| S6C: PPR (50Hz) | Students t test, unpaired, two-tailed | t=0.9279, df=19 | *p=0.3651* | WT n=12, $CK1\delta_{T44A}$ n=11 | No. of neurons (1 neuron per acute slice) |
| S6D: Cumulative IPSC amplitude (10Hz) | two-way ANOVA | Stim x Genotype: F (29, 290) = 0.07050, p>0.9999 Stim: F (1.021, 10.21) = 29.41, p=0.0003 Genotype: F (1, 10) = 0.01786, p=0.8963 | *p>0.9999* | WT n=11, $CK1\delta_{T44A}$ n=11 No. of neurons (1 neuron per acute slice) | |
| | ANCOVA (y-intercepts) | F (1, 272) = 0.8894 | *p=0.3465* | | |
| S6E: Cumulative IPSC amplitude (20Hz) | two-way ANOVA | Stim x Genotype: F (29, 435) = 0.1928, p>0.9999 Stim: F (1.119, 16.78) = 42.04, p<0.0001 | *p>0.9999* | WT n=11, $CK1\delta_{T44A}$ n=10 | No. of neurons (1 neuron per acute slice) |

| | | | | | |
|---|---|---|---|---|---|
| | | Genotype: F (1, 15) = 0.4734, p=0.5019 | | | |
| | ANCOVA (y-intercepts) | F (1, 249) = 0.05291 | *p=0.8183* | | |
| S6F: Cumulative IPSC amplitude (50Hz) | two-way ANOVA | Stim x Genotype: F (29, 377) = 0.4718, p=0.9918 Stim: F (1.081, 14.05) = 29.45, p<0.0001 Genotype: F (1, 13) = 0.07103, p=0.7940 | *p=0.9918* | WT n=11, CK1$\delta_{T44A}$ n=11 | No. of neurons (1 neuron per acute slice) |
| | ANCOVA (y-intercepts) | F (1, 261) = 0.3670 | *p=0.5450* | | |
| S6G: y-intercepts and slopes (50Hz) | Students t test, unpaired, two-tailed | t=0.5922, df=20 | *p=0.5604* (y-intercept) | WT n=11, CK1$\delta_{T44A}$ n=11 | No. of neurons (1 neuron per acute slice) |
| | | t=0.7728, df=20 | *p=0.4385* (Slope) | | |
| S6H: y-intercepts and slopes (20Hz) | Students t test, unpaired, two-tailed | t=0.7139, df=19 | *p=0.4840* (y-intercept) | WT n=11, CK1$\delta_{T44A}$ n=10 | No. of neurons (1 neuron per acute slice) |
| | | u=37 | *p=0.2230* (Slope) | | |
| S6I: y-intercepts and slopes (10Hz) | Students t test, unpaired, two-tailed | t=0.7496, df=19 | *p=0.4627* (y-intercept) | WT n=11, CK1$\delta_{T44A}$ n=11 | No. of neurons (1 neuron per acute slice) |
| | | t=0.8842, df=19 | *p=0.3876* (Slope) | | |
| **Figure S7** | | | | | |

| | | | | | |
|---|---|---|---|---|---|
| S7B: input-output relationship | Two-way repeated measures ANOVA, | Row x Column: F (9, 54) = 3.976, *P=0.0006* Row: F (9, 54) = 6.547, *P<0.0001* Column: F (1, 6) = 8.146, *P=0.0290* | *p<0.001* | WT n=4, CK1δ$_{T44A}$ n=4 | No. of neurons (1 neuron per acute slice) |
| **Figure S8** | | | | | |
| S8C: Normalized EPSC amplitude (10Hz) | Two-way repeated measures ANOVA, | Stim x Genotype: F (9, 99) = 1.104, *p=0.3672* Stim: F (9, 99) = 24.30, *p<0.0001* Genotype: F (1, 11) = 0.9110, p=0.3603 | *p=0.1391* | WT n=10, CK1δ$_{T44A}$ n=13 | No. of neurons (1 neuron per acute slice) |
| S8C: PPR (10Hz) | Students t test, unpaired, two-tailed | t=0.2328, df=11 | *p=0.3651* | WT n=10, CK1δ$_{T44A}$ n=13 | No. of neurons (1 neuron per acute slice) |
| S8D: Normalized EPSC amplitude (20Hz) | Two-way repeated measures ANOVA, | Stim x Genotype: F (9, 207) = 2.556, *p=0.0083* Stim: F (9, 207) = 18.34, *p<0.0001* Genotype: F (1, 23) = 0.7615, *p=0.3919* | *p<0.01* | WT n=10, CK1δ$_{T44A}$ n=13 | No. of neurons (1 neuron per acute slice) |

| | | | | | |
|---|---|---|---|---|---|
| S8D: PPR (20Hz) | Students t test, unpaired, two-tailed | t=0.8510, df=23 | *p=0.4035* | WT n=10, CK1$\delta_{T44A}$ n=13 | No. of neurons (1 neuron per acute slice) |
| S8E: Normalized EPSC amplitude (50Hz) | Two-way repeated measures ANOVA, | Stim x Genotype: F (9, 198) = 2.530, *p=0.0091* Stim: F (9, 198) = 13.45, *p<0.0001* Genotype: F (1, 22) = 0.2013, *p=0.6581* | *p<0.01* | WT n=10, CK1$\delta_{T44A}$ n=13 | No. of neurons (1 neuron per acute slice) |
| S8E: PPR (50Hz) | Students t test, unpaired, two-tailed | t=0.4070, df=21 | *p=0.6881* | WT n=10, CK1$\delta_{T44A}$ n=13 | No. of neurons (1 neuron per acute slice) |
| S8F: Normalized IPSC amplitude (10Hz) | Two-way repeated measures ANOVA, | Stim x Genotype: F (9, 108) = 0.7472, *p=0.6650* Stim: F (9, 108) = 30.50, *p<0.0001* Genotype: F (1, 12) = 0.002288, *p=0.9626* | *p=0.6650* | WT n=9, CK1$\delta_{T44A}$ n=15 | No. of neurons (1 neuron per acute slice) |
| S8F: PPR (10Hz) | Students t test, unpaired, two-tailed | t=0.3306, df=12 | *p=0.7466* | WT n=9, CK1$\delta_{T44A}$ n=15 | No. of neurons (1 neuron per acute slice) |
| S8G: Normalized IPSC | Two-way repeated | Stim x Genotype: F (9, 207) = 0.4249, *p=0.9208* | *p=0.9208* | WT n=9, CK1$\delta_{T44A}$ n=15 | No. of neurons (1 |

| | | | | | |
|---|---|---|---|---|---|
| amplitude (20Hz) | measures ANOVA, | Stim: F (9, 207) = 36.41, p<0.0001 Genotype: F (1, 23) = 0.001561, p=0.9688 | | | neuron per acute slice) |
| S8G: PPR (20Hz) | Students t test, unpaired, two-tailed | t=0.4194, df=23 | *p=0.6786* | WT n=9, CK1$\delta_{T44A}$ n=15 | No. of neurons (1 neuron per acute slice) |
| S8H: Normalized IPSC amplitude (50Hz) | Two-way repeated measures ANOVA, | Stim x Genotype: F (9, 189) = 0.6929, *p=0.7149* Stim: F (9, 189) = 19.57, *p<0.0001* Genotype: F (1, 21) = 0.0004248, *p=0.9838* | *p=0.7149* | WT n=9, CK1$\delta_{T44A}$ n=15 | No. of neurons (1 neuron per acute slice) |
| S8H: PPR (50Hz) | Students t test, unpaired, two-tailed | t=0.4512, df=22 | *p=0.6562* | WT n=9, CK1$\delta_{T44A}$ n=15 | No. of neurons (1 neuron per acute slice) |
| **Figure S9** | | | | | |
| S9B: SD velocity | Two-way ANOVA, Tukey's test for multiple comparisons | Interactions: WT control vs CK1$\delta_{T44A}$ control: *p=0.5881* WT control vs WT low $Ca^{2+}$: *p=0.0611* | Genotype *p=0.5087*, Treatment *p=0.0132* | Control (WT, CK1$\delta_{T44A}$ n=23) Low $Ca^{2+}$ ACSF (WT, CK1$\delta_{T44A}$ n=20) | No. of slices |

| | | | | | |
|---|---|---|---|---|---|
| | | $CK1\delta_{T44A}$ control vs $CK1\delta_{T44A}$ low $Ca^{2+}$: *p=0.726*<br>WT low $Ca^{2+}$ vs $CK1\delta_{T44A}$ low $Ca^{2+}$: *p=0.993* | | | |
| S9C: SD propagation | Two-way ANOVA, Tukey's test for multiple comparisons | Interactions:<br>WT control vs $CK1\delta_{T44A}$ control: *p=0.995*<br>WT control vs WT low $Ca^{2+}$: *p<0.0001*<br>$CK1\delta_{T44A}$ control vs $CK1\delta_{T44A}$ low $Ca^{2+}$: *p<0.0001*<br>WT low $Ca^{2+}$ vs $CK1\delta_{T44A}$ low $Ca^{2+}$: p=0.4415 | Genotype *p=0.5087,* Treatment *p<0.0001* | Control (WT, $CK1\delta_{T44A}$ n=23)<br>Low $Ca^{2+}$ ACSF (WT, $CK1\delta_{T44A}$ n=20) | No. of slices |

#### Table S2. Description of the key resources

| **Reagents or resources** | **Source** | **Identifier** |
|---|---|---|
| **Viral strain** | | |
| AAV1.hSyn.iGluSnFr.WPRE.SV40 | UPenn Vector Core | AV-1-PV2723 |
| **Experimental models** | | |
| Human CK1δ mutant ($CKI\delta_{T44A}$) BAC transgenic mice | Xu et. Al., 2005 | N/A |
| **Chemical and other reagents** | | |
| Tetrodotoxin (TTX) | Tocris | Catalog #: 1078 |
| DL-2-Amino-5-phosphonopentanoic acid (DL-APV) | Tocris | Catalog #: 0105 |
| 6,7-Dinitroquinoxaline-2,3-dione disodium salt (DNQX) | Tocris | Catalog #: 2312 |
| MK-801 maleate | Tocris | Catalog #: 0924 |
| Picrotoxin (PTX) | Tocris | Catalog #: 1128 |
| QX-314 bromide | Tocris | Catalog #: 1014 |
| **Software and algorithms** | | |
| MATLAB 2017a | Mathworks | https://www.mathworks.com/ |
| Prism 8 | Graphpad | https://www.graphpad.com/ |
| Fiji | ImageJ | https://imagej.net/Welcome |
